## Extended Data and Supplementary Information for "Thermodynamic dissipation constrains metabolic versatility of unicellular growth"

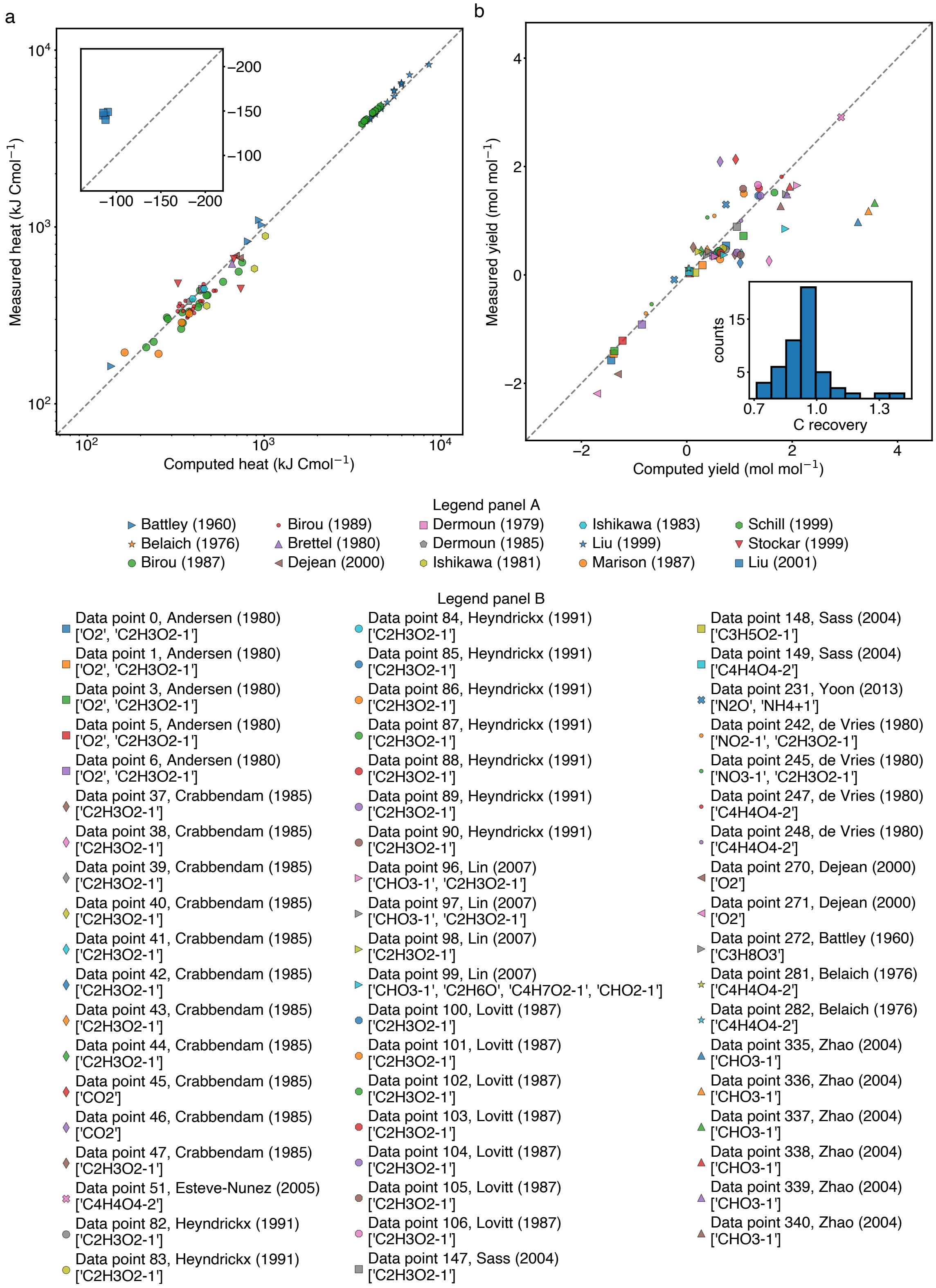


Extended Data Fig.1. **Validation of the first law of thermodynamics and element conservation. a.** (Minus) the heat per Cmol of biomass predicted using enthalpy balance is plotted against direct calorimetric measurements. The legend, below, includes the bibliographic reference (see main text for labelling by types). The inset corresponds to endothermic measurements. The deviations from the dashed line, which is not a fit, are small. This constitutes a validation of our approach. **b.** Measured yields of substrates and products, reported in the file `measured_yields.csv`, are plotted against the corresponding yields predicted using element conservation, reported in the file `yields.csv`. The sign distinguishes substrates, negative in this plot, from products, positive. The yields are reported in moles per moles of electron donor. The legend below indicates the "entry" number of the experiment where the fluxes were measured, the bibliographic reference of the experiment, and the stoichiometric formula of the yields measured, all of the above consistently with the notation used in the data-files. Values of carbon recovery, parsed from the primary sources used in this work, are plotted as an inset. As in the case of heat, predictions match measurements closely, which validates our usage of element conservation.

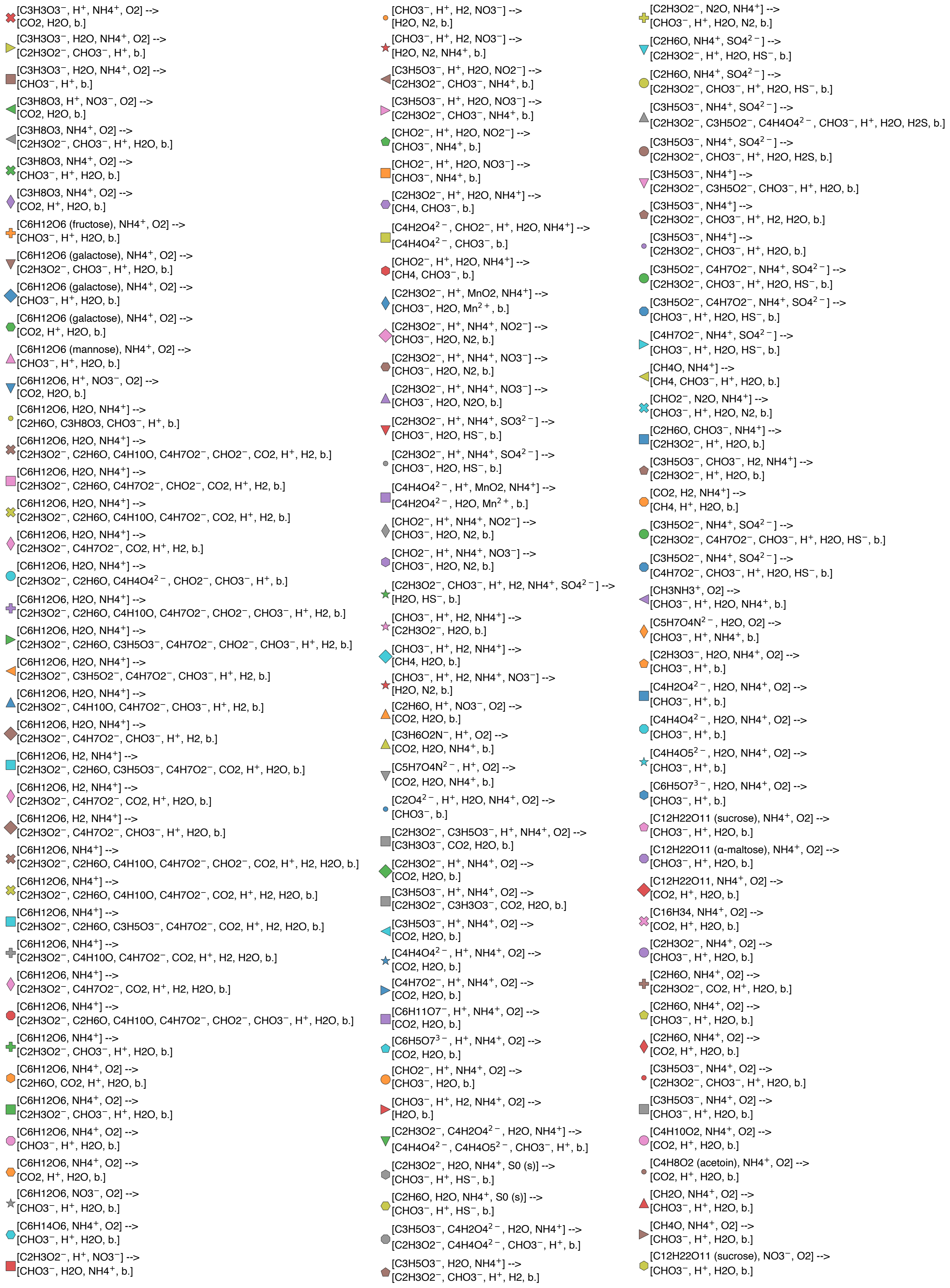


Extended Data Fig.2. **Legend of all metabolic types in the dataset.** This is a comprehensive list of all metabolic types in the dataset. Each symbol/color denotes the macrochemical equation next to it, written as `[substrates]-->[products]`, with the letter `b` denoting biomass. This legend is later used in Extended Data Fig.5, Extended Data Fig.6 and Extended Data Fig.9

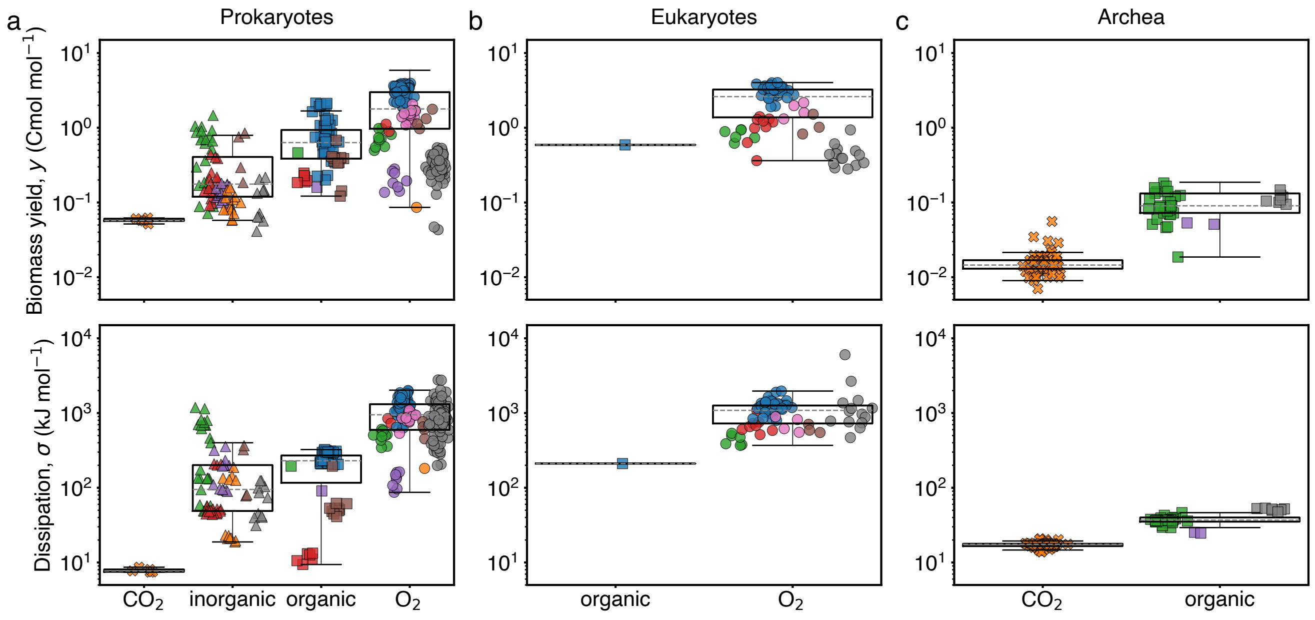


Extended Data Fig.3. **Yield and dissipation separated by domains of life. a-c.** Biomass yield variability (upper row) as in Fig 2a, and dissipation variability (lower row) as in Fig 2b, for prokaryotes (a), eukaryotes (b), and archea (c). While eukaryotes have typically higher yield and dissipation than prokaryotes, the values are comparable to those of prokaryotes with similar metabolic types (e.g. aerobic respiratory types). While archae have generally lower yield and dissipation than prokaryotes, there are many prokaryotic data with comparably low yields. Moreover, autotrophic methanogenesis type is found exclusively in archea, and is the reponsible for the lowest yields and dissipations in the domain. Overall, we can not ascribe the observed variability to differences in domains of life.

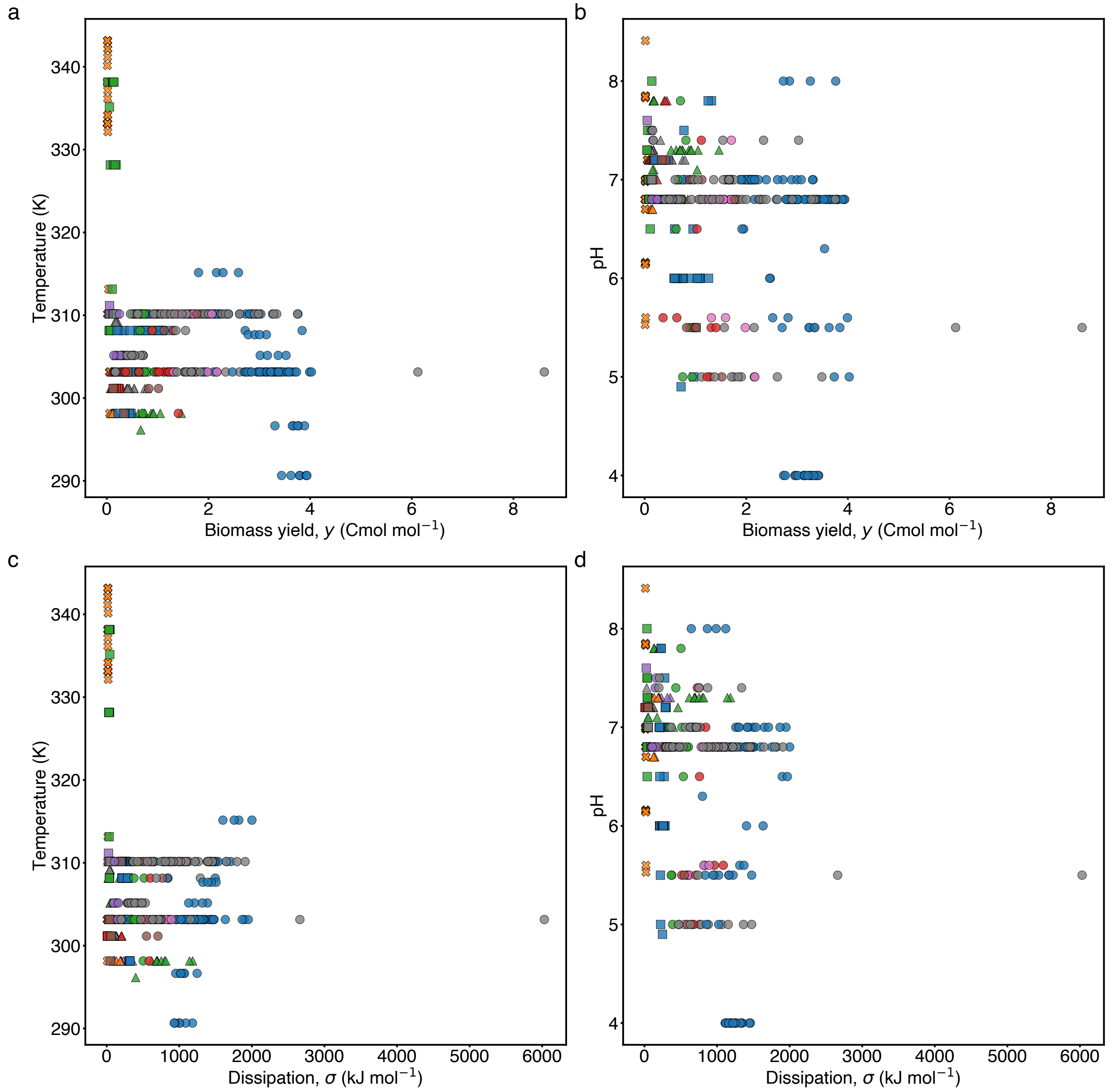


Extended Data Fig.4. **Dissipation and yield as functions of temperature and pH. a.** Scatter plot of yield against temperature at which the corresponding experiment was performed. **b.** Scatter plot of yield vs pH of the media. **c.** Scatter plot of dissipation against temperature. **d.** Scatter plot of dissipation against pH. The extent of temperatures explored is large, $\approx 50^{\circ}$C, as also is the range of pH conditions, $\approx 4$. Despite this, no consistent relationship between yield and temperature or pH is observed. The same is true for dissipation, despite the fact that pH and temperature corrections were included in computing the dissipation, see SI G.

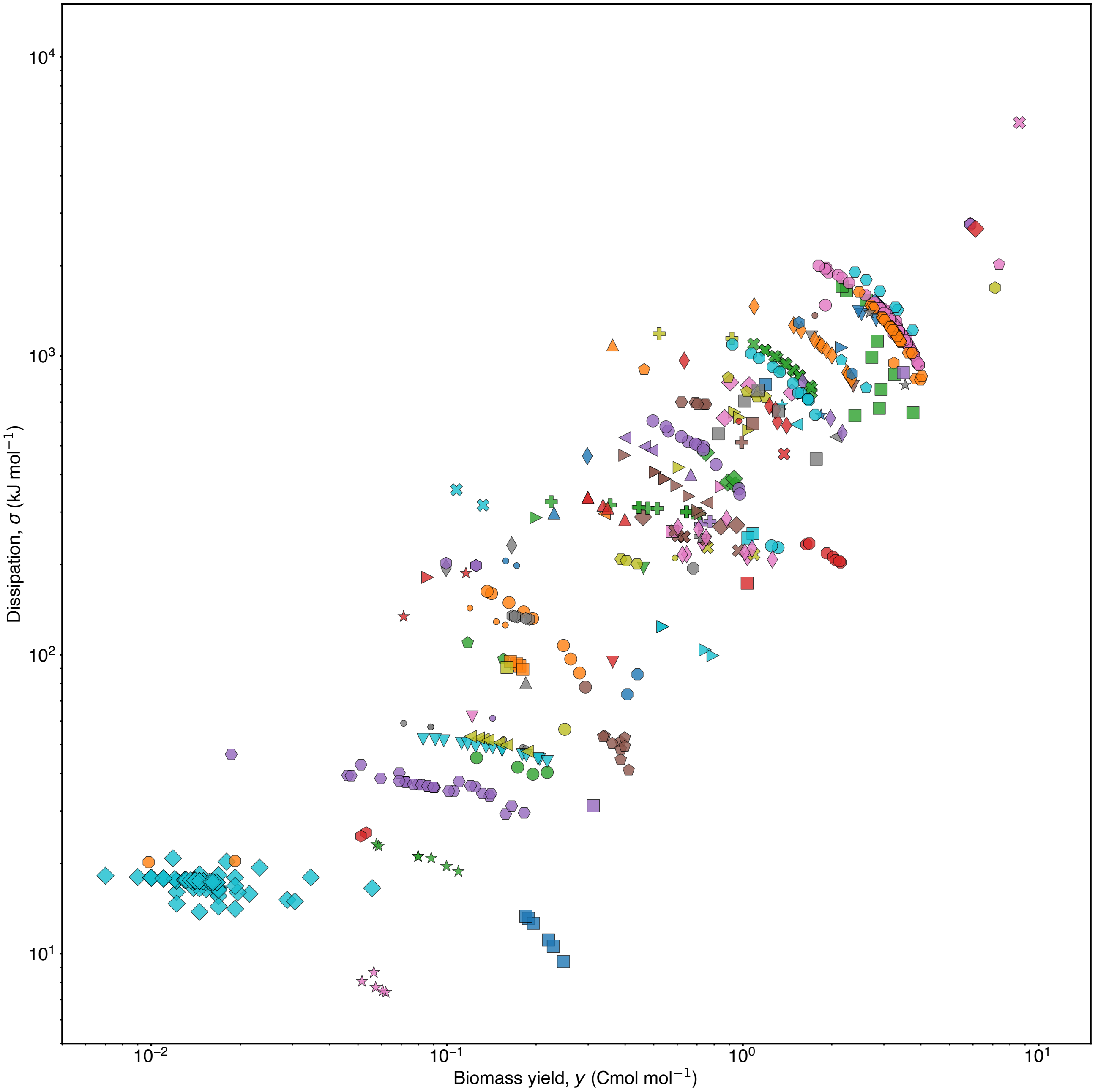


Extended Data Fig.5. **Yield-dissipation relation for the complete dataset.** Analogue of Fig 3a using the comprehensive labeling from Extended Data Fig.2, which distinguishes all different metabolic types included in the dataset.

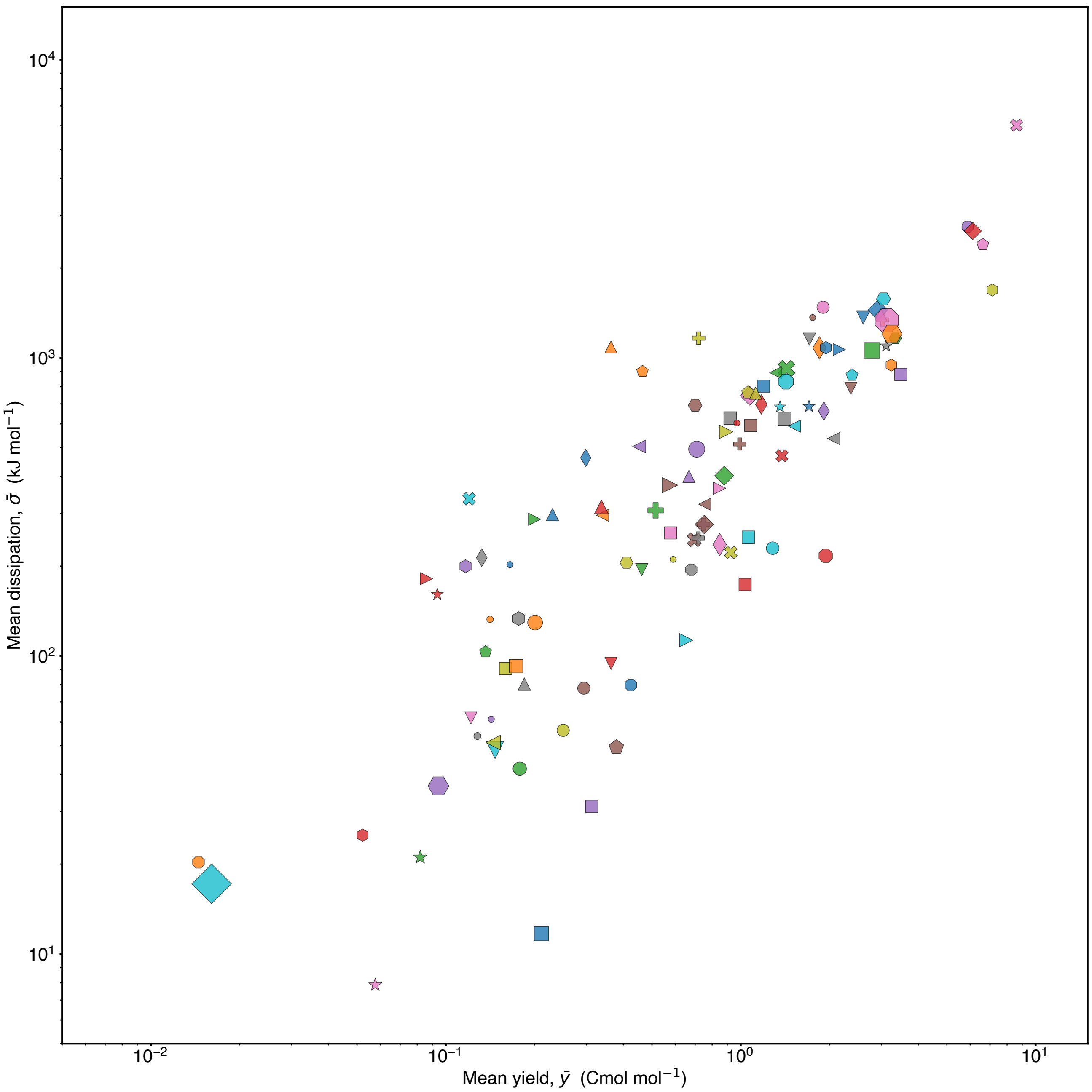


Extended Data Fig.6. **Yield-dissipation relation across metabolic types.** Analogue of Fig 3b using the comprehensive labeling from Extended Data Fig.2, which distinguishes all different metabolic types included in the dataset. The size of the marker is proportional to the number of data points.

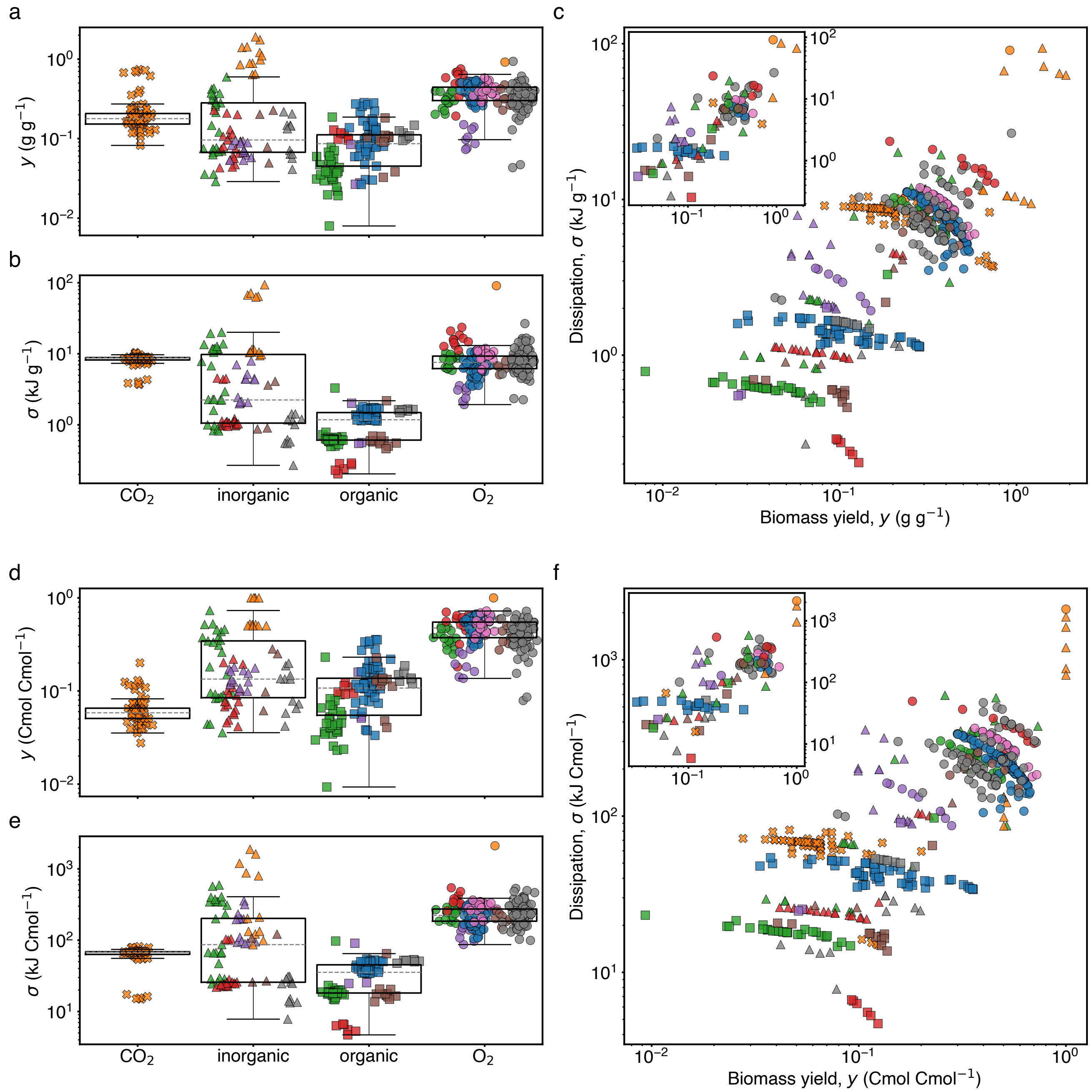


Extended Data Fig.7. **Dissipation, yield and their relationship for two alternative choices of units. a** and **b.** Analogue of Fig. 2 using units of gram per gram for the biomass yield and kJ per gram for the dissipation. Note that the range of the data is somewhat reduced, but still spans over two orders of magnitude for yield, and three for dissipation. **c.** Analogue of Fig. 3a repeated for the units of panels a and b, which clearly preserves the correlation between dissipation and yield. Note that the data using $H_2$ as electron donor has substantially moved within the plot, due to the low weight of this substrate. **c inset.** Analogue of Fig. 3b, mean dissipation $\bar{\sigma}$ plotted against mean yield $\bar{y}$, for each type using the units of panel c. **d**, **e** and **f.** The same as panels a-c but for units of Cmol per Cmol, with similar conclusions. Note that the data using $H_2$ as electron donor is now expressed per carbon mole of carbon source ($CO_2$) instead.

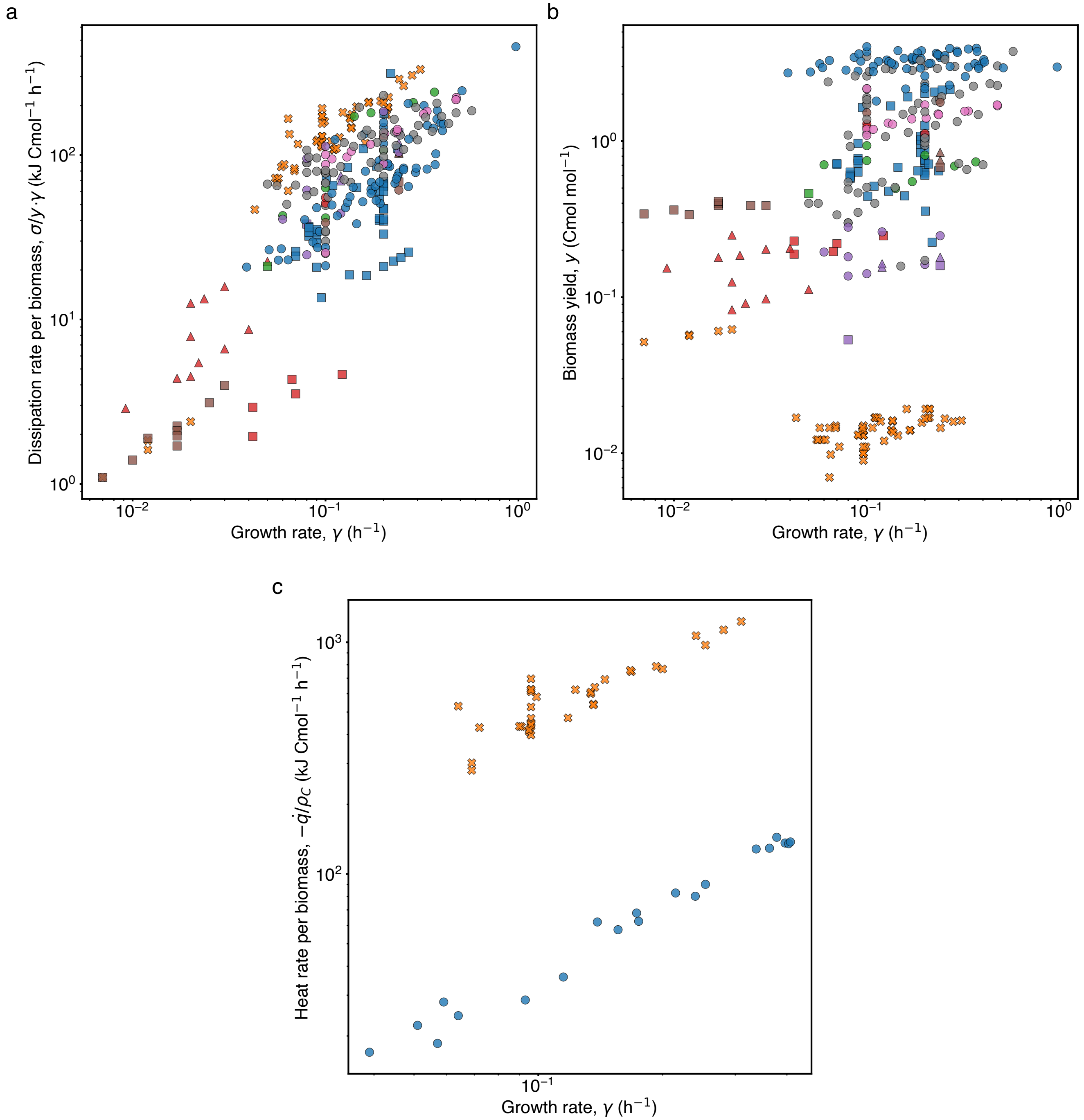


Extended Data Fig.8. **Characterization of dissipation per unit time for chemostated data. a.** Dissipation per unit biomass and time, $(\sigma/y)\gamma$, plotted against growth rate. The median power dissipated per biomass grown is 81 kJ $\mathrm{Cmol}^{-1}\,\mathrm{h}^{-1}$, or 0.9 W $\mathrm{g}^{-1}$. We find a roughly linear scaling for about two decades of growth rate. This is in agreement with the notion of cost conservation, which is robust to changes in growth rate, and compatible with observations in Niebel et al. [1]. **b.** Biomass yield plotted against growth rate. We do not find any clear relation (i.e. no yield-growth rate trade-off), not within metabolic type nor across metabolic types. **c.** Calorimetric measurements of heat produced per unit biomass and time, $-\dot{q}/\rho_{\mathrm{C}}$, plotted against growth rate. Autotrophic methanogenesis, orange crosses, shows a noticeable difference between heat and dissipation (panel a). This is due to the big contribution of chemical entropy to the dissipation, Eq. (B17). Data are from [2–5].

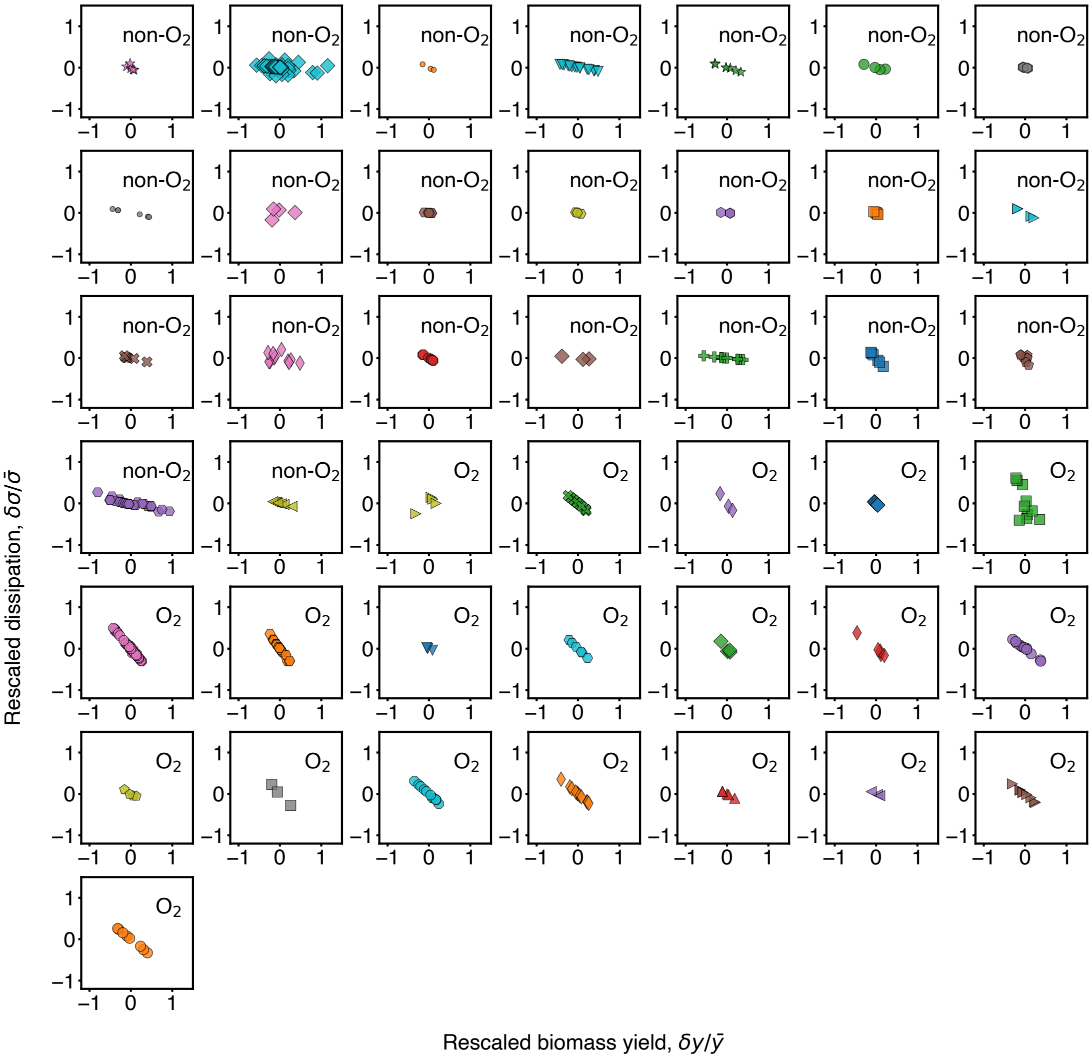


Extended Data Fig.9. **Examples of the relationship between deviations in dissipation and yield of different metabolic types.** Analogue of Fig. 3c, dedicating an individual panel to each metabolic type with three or more data points. On the x-axis, the normalized deviation in yield, and on the y-axis, the normalized deviation in dissipation. The legend is that in Extended Data Fig.2.

### Supplementary Information

#### CONTENTS

A. Summary of supplementary files 12

B. Non-equilibrium thermodynamics of unicellular growth 13
1. Mass conservation and steady balanced growth 13
2. Definition of yield 14
3. Element conservation and macrochemical equation 16
4. Thermodynamics 16

C. Defining the cost across and the coefficient within metabolic types 18

D. Alternative definitions for the thermodynamic cost of growth 19

E. The origin of the cost across types, $\alpha$, and of the within type coefficient, $\beta$ 22
1. Analytical relation between yield and dissipation 22
2. Variations of yield and dissipation across and within metabolic type 22
3. Efficiency 24
4. Origin of the differences in deviation coefficient $\beta$ for aerobic and anaerobic types 24

F. A thermodynamic database of microbial growth 25
1. Data curation and validation from primary literature sources 25
2. Units and typical values of variables used 26
3. Comments on data curation and parsing from primary literature 26

G. Thermodynamic properties of substrates and products 32

H. Thermodynamic properties of biomass 34
1. Biomass composition 34
2. Enthalpy, entropy and free energy of biomass 34

I. Dataset validation 36

J. Symbols 37

#### Appendix A: Summary of supplementary files

The results of this work are supported by multiple data-files appended to it, which we describe below.

- measured_yields.csv: This file contains the experimentally measured yields that we parsed from the primary sources. Each row corresponds to an individual entry, i.e. an experiment where one or more yields were measured. We identify the entries by a number, denoted "entry", which is consistently used across all the data-files. Each column corresponds to a chemical species, denoted by its stoichiometric formula. The formula is used consistently throughout the data-files and in the file thermo_chem.csv we also report the name of the chemical species. We report the values of the yields as moles per moles of electron donor (carbon moles in the case of biomass). When the primary source reported the yields in a different form, e.g. in different units or as individual fluxes, we applied the transformation needed. In this data-file, the sign is negative for the chemical species consumed and positive for the ones produced.

- yields.csv: This file contains all the yields of biomass and chemical species exchanged as metabolic substrates and products. The rows (entries) contain the yields of the corresponding experiment, whose values were either measured or computed as explained below. Each column corresponds to a chemical species identified by its stoichiometric formula. For each entry, we enforced element conservation, which provides a set of equations for the yields, SI B 3 and I. To solve these equations, we used the measured yield of biomass as input. When the system of equations needed multiple yields to be determined, we chose these extra yields among the measured yields that we parsed. The remaining yields were computed solving the equations for the balance of elements. In some occasions, we could parse more measured yields than those needed to determine the system of equations. We decided which yields to use to determine each system on the basis of the measurements methods reported in the original reference, trying to select the more reliable ones. The yields chosen are reported as "extra yields used". The extra yields not used to determine the system were used to validate the dataset, SI I and Fig. 1a. In all cases, the yield of chemical species is computed relative to the electron donor, and reported in units of moles per moles of electron donor. In this data-file, the sign is negative for the chemical species consumed and positive for the ones produced. For example, the yield of electron donor trivially takes the value $-1$, whereas the yield of biomass is always a positive number. Finally, the file contains the biomass stoichiometry of each entry, used to compute the element balance.

- thermo_growth_dat.csv: This file contains the auxiliary information for each entry, including microbial species, temperature, pH, growth rate, a list of chemical products and substrates of metabolism, heat measured and carbon balance (when available). This information was retrieved from the original sources of the data, with possibly the exception of the products. In some cases indeed, we completed the list of metabolic products beyond what explicitly stated in the papers, on the basis of well known metabolic pathways (e.g. if glucose was metabolized respiring oxygen, we included carbon dioxide as product even if not mentioned).

- biomass_composition.csv: This file contains data on the stoichiometric composition of biomass. Each row refers to an organism identified by the name of the species and optionally the strain or additional qualifiers. We reported the hydrogen, oxygen and nitrogen content of biomass normalized by carbon content. We discarded other elements when (very rarely) reported. The phylum and domain of the organism, as well as the reference where the stoichiometric data was taken from, were also reported.

- thermo_chem.csv: This file contains the thermodynamic properties of all chemical species in the dataset. Each row corresponds to a chemical species, identified by its stoichiometric formula, consistently with the other files, and its name. The free energies of formation are reported in standard conditions, i.e. 1 M concentration at 298 Kelvin. We further reported the enthalpies of formation, used to validate the data that provided calorimetric measurements and to correct the use of standard free energies at non-standard temperature, according to the Gibbs-Helmholtz equation, Eq. (G2).

- thermo_growth_computed.csv This file contains the main thermodynamic quantities computed (excluding biomass yield) as described in the text and SI, including dissipation $\sigma$ (free energy dissipated per electron donor consumed), free energy dissipated per biomass produced $\sigma/y$, anabolic force $r_{\rm b}$, catabolic force $r_{\rm ed}$, thermodynamic efficiency $\eta$, biomass (molecular) weight $m_{\rm b}$,free energy and enthalpy of formation of biomass $g/\rho_{\rm C}$ and $h/\rho_{\rm C}$. In this file, we associated a name to each metabolic type, providing a coarser grouping than the macrochemical equation. For example, "anaerobic fermentation of glucose" does not distinguish the different fermentation products. We remark that a metabolic type is defined by the set of substrates and products (i.e. macrochemical equation), but with no distinction between (de)protonated forms of chemical species (e.g. we considered equivalent acetic acid and acetate, ignoring charges and protons) nor between carbon dioxide and bicarbonate.

#### Appendix B: Non-equilibrium thermodynamics of unicellular growth

##### 1. Mass conservation and steady balanced growth

We describe a cellular colony as a dilute solution of $P$ biomass constituents, $i = 1, \dots, P$, at abundances $N_i$. The constituents span any cellular component, including metabolites, nucleic acids and proteins, and are characterized by their mass $m_i$. The mass abundances are thus defined as $M_i = m_i N_i$ and the total biomass as $M = \sum_i M_i$. This biomass occupies a volume $V$ at constant pressure $\Pi$ (not to be confused with the volume of the vessel where the colony grows, $V^{\mathrm{v}}$, the two related through the volume fraction $\phi \equiv V/V^{\mathrm{v}}$), which allows to define the dry biomass density $\rho = M/V$ (i.e. weight of dry mass per volume of biomass).

To sustain steady growth, cells import substrates from the surrounding media into the colony at fluxes $F_i^+ > 0$ and export products at fluxes $-F_i^- < 0$. Furthermore, inside the cell matter is transformed through reaction fluxes $K_j$ spanning chemical reactions $j = 1, \dots, R$. This network of chemical reactions is characterized by the stoichiometric matrix $a_{ij}$. Within this framework, mass conservation for each constituent is captured by

$$\dot{N}_i = F_i^+ - F_i^- + \sum_j a_{ij} K_j \quad . \tag{B1}$$

Notice that for each chemical $i$, either $F_i^+ = 0$ if $i$ is a product ($i \in \mathrm{pd}$), $F_i^- = 0$ if it is a substrate ($i \in \mathrm{sb}$), or $F_i^+ = F_i^- = 0$ for all the other biomass constituents.

The growth rate of the colony is the amount of biomass produced per unit biomass, and so for a closed vessel we have $\gamma \equiv \dot{M}/M$. Similarly, the volume of the colony, $V$, expands as $\dot{V} = \gamma V$. Taking this into consideration, we can write an intensive form of Eq. (B1) by defining the molar concentrations of chemicals inside the colony as $c_i = N_i/V$ (we similarly define densities as $\rho_i = m_i c_i$, with $\rho = \sum_i \rho_i$). Noting that $\dot{c}_i = \dot{N}_i/V - \gamma c_i$, we have

$$\dot{c}_i = f_i^+ - f_i^- - \gamma c_i + \sum_j a_{ij} \nu_j \quad , \tag{B2}$$

where we have defined the molar import/export flux densities as $f_i^{+/-} = F_i^{+/-}/V$, and the molar reaction flux densities in chemical reactions as $\nu_j = K_j/V$. In this work, we only consider the case of steady balanced growth, in which the growth rate $\gamma$ is time independent and equal for all constituents, i.e. $\dot{N}_i = \gamma N_i$. This corresponds to exponential growth conditions in which the composition of the cells is not changing. In that case, we have that $\dot{c}_i = 0$, and Eq. (B2) reduces to

$$\gamma c_i = f_i^+ - f_i^- + \sum_j a_{ij} \nu_j \quad . \tag{B3}$$

Since the fluxes $F_i^{+/-}$ and $K_j$ scale as $\sim N_i$, the quantities $f_i^{+/-}$ and $\nu_j$ are intensive and time-independent. We note that this description of microbial growth is analogous to that which forms the basis of Flux Balance Analysis Varma and Palsson [6].

In a batch culture, steady balanced growth is achieved during the exponential stage of cell growth in a vessel, i.e. when $\phi \ll 1$, prior to the cellular colony reaching carrying capacity. Steady balanced growth can also be achieved in a chemostat. To see this, note that in a chemostat an amount of volume is constantly added and removed per unit time, $G$, with the dilution rate defined as $d = G/V^{\mathrm{v}}$. The volume added contains nutrients, whereas the volume removed contains all constituents present in the vessel, including biomass. The balance of biomass is, therefore, $\dot{M} = \gamma M - M d$, which reaches a steady state for $d = \gamma$. Equation (B1) is also modified to

$$\dot{N}_i = F_i^+ - F_i^- + \sum_j a_{ij} K_j - N_i d \quad , \tag{B4}$$

and a steady state is reached when the abundances of cellular components are constant within the vessel, i.e. $\dot{N}_i = 0$. Applying this condition in Eq. (B4), identifying $d = \gamma$, and dividing by the volume of the colony $V$ results in Eq. (B3). Therefore, the equation of steady balanced growth applies equally to chemostats at steady state and batch cultures during exponential growth. Note that in writing Eq. (B4) we took into consideration that the exchange fluxes $F_i^{+/-}$ refer to the cellular colony, and not to the vessel.

We emphasize that the concentration $c_i$ represents the amount of constituent $i$ per volume of the cellular colony. Alternatively, one can define concentrations per dry-mass weight of cellular colony, the difference between two being a rescaling by the density $\rho$. Since our results are presented as ratios of per colony volume quantities, this choice is arbitrary. We further discuss this point and other unit choices in section F 2.

##### 2. Definition of yield

The biomass yield is an important physiological quantity that characterizes growth and plays a central role in our work. Broadly speaking, the growth yield or biomass yield, $y$, quantifies the amount of biomass grown per unit of nutrient consumed. We now review the precise definition of growth yield, and how it is inferred for two different experimental scenarios: chemostated cultures and batch cultures.

We first start with the case of a chemostated culture. First, we write the kinetics of biomass and nutrients in the chemostat. Mass conservation in a chemostat is expressed through Eq. (B4). Defining the exchange mass fluxes $\mathcal{M}_i^{+/-} = m_i F_i^{+/-}$, as well as the net exchange mass fluxes $\mathcal{M}^{+/-} = \sum_i \mathcal{M}_i^{+/-}$, leads to

$$\dot{M} = \mathcal{M}^+ - \mathcal{M}^- - dM \quad , \tag{B5}$$

where we can identify the growth rate as $\gamma = (\mathcal{M}^+ - \mathcal{M}^-)/M$. For the case of any chemical $i$ consumed or produced by the cell $i$, and in particular for the electron donor $i$ = ed, we have that the total abundance in the vessel is composed of the $N_{\mathrm{ed}}$ molecules inside the colony plus the $N'_{\mathrm{ed}}$ diluted in the surrounding media. Mass conservation for the abundances in the media is therefore given by

$$\dot{M}'_{\mathrm{ed}} = \rho^{\mathrm{in}}_{\mathrm{ed}} G - \rho'_{\mathrm{ed}} G - \mathcal{M}^+_{\mathrm{ed}} \quad , \tag{B6}$$

where $\rho^{\mathrm{in}}_{\mathrm{ed}}$ is the density of electron donor in the media that is flowed into the chemostat, and we have defined the mass of electron donor outside the colony $M'_{\mathrm{ed}} = m_{\mathrm{ed}} N'_{\mathrm{ed}}$ as well as the density outside the colony $\rho'_{\mathrm{ed}} = M'_{\mathrm{ed}}/V^{\mathrm{v}}$, now relative to the volume of the vessel. A steady state for a chemostat is achieved for $d = \gamma$ and a concentration of electron donor in the vessel $\rho'_{\mathrm{ed}} = \rho^{\mathrm{in}}_{\mathrm{ed}} - \mathcal{M}^+_{\mathrm{ed}}/G$.

In this chemostat setting the yield is defined as

$$Y \equiv \frac{\gamma M}{\mathcal{M}^+_{\mathrm{ed}}} \quad , \tag{B7}$$

which is the expression commonly used in the primary sources of our study. Note that at steady state we have $\mathcal{M}^+_{\mathrm{ed}} = G(\rho^{\mathrm{in}}_{\mathrm{ed}} - \rho'_{\mathrm{ed}})$, which allows to determine the denominator directly from measuring the concentration in outflux from the chemostat. Often, nutrients are limiting in the media and so most nutrients are depleted, $\rho'_{\mathrm{ed}} \ll \rho^{\mathrm{in}}_{\mathrm{ed}}$. In this case, we can approximate $\mathcal{M}^+_{\mathrm{ed}} \approx G\rho^{\mathrm{in}}_{\mathrm{ed}}$. Note also that the stability of the steady state requires further assumptions on the dependence of the growth rate on the electron donor abundance, e.g. Monod law $\gamma = \gamma_{\max} M'_{\mathrm{ed}}/(K + M'_{\mathrm{ed}})$, with $K$ an effective constant and $\gamma_{\max}$ the maximal growth rate Monod [7]. The definition of yield allows to write the chemostat equations in the more familiar form

$$\dot{M} = \gamma M - dM \tag{B8}$$

$$\dot{M}'_{\mathrm{ed}} = \rho^{\mathrm{in}}_{\mathrm{ed}} G - \rho'_{\mathrm{ed}} G - \gamma M/Y \quad . \tag{B9}$$

The expression of yield in Eq. (B7) is written using extensive quantities, but it is straightforward to divide by the cellular volume and obtain

$$Y = \frac{\gamma \rho}{m_{\mathrm{ed}} f^+_{\mathrm{ed}}} \quad . \tag{B10}$$

Furthermore, while the yield is a dimensionless quantity, it was calculated as a ratio of mass fluxes. Instead, we can define an expression of the yield as a ratio of carbon molar flux for biomass, and molar flux for electron donor. To achieve this, we simply multiply the above yield by the conversion factor $m_{\mathrm{ed}}/m_{\mathrm{b}}$, with $m_{\mathrm{b}}$ the molecular weight of biomass, see also F 2. The resulting form for the yield is then

$$y = \frac{\gamma \rho_{\mathrm{C}}}{f^+_{\mathrm{ed}}} \quad , \tag{B11}$$

with $\rho_{\mathrm{C}} = \rho/m_{\mathrm{b}}$ the dry biomass density in carbon moles. This is the expression that was used in the main text.

We now discuss the case of *batch culture*. Conservation for the biomass of the colony and the mass of electron donor outside the colony is simply described by

$$\dot{M} = \mathcal{M}^+ - \mathcal{M}^- \tag{B12}$$

$$\dot{M}'_{\mathrm{ed}} = -\mathcal{M}^+_{\mathrm{ed}} \quad , \tag{B13}$$

where as before the growth rate is given by $\gamma = (\mathcal{M}^{+} - \mathcal{M}^{-})/M$. At early stages of growth the electron donor is in great abundance, and the growth rate can be considered independent of $M'_{\rm ed}$ (for Monod's law we have $\gamma \approx \gamma_{\rm max}$). In this case, we can use the definition of yield in Eq. (B7) to obtain the time-dependent solutions $M(t) = M(0)\exp(\gamma t)$ and $M'_{\rm ed} = M'_{\rm ed}(0) - (M(t) - M(0))/Y$. From this, we obtain that during the exponential phase the yield can also be written as

$$Y = -\frac{M(t) - M(0)}{M'_{\rm ed} - M'_{\rm ed}(0)} \quad , \tag{B14}$$

which is the expression used for the batch culture experiments that we analyzed (transforming to $y$ is, as before, straightforward). This expression is strictly correct for the case in which the experiment was arrested during exponential growth (note that the lag-phase, in which nutrient is not depleted and biomass does not grow, does not affect this result). However, Eq. (B14) can also be used as an approximation for the yield when the experiment is run until depletion of substrate. The accuracy of this approximation then depends on the duration of the transition from exponential growth to saturation.

We note that, besides the biomass yield, the yield of any chemical can be computed, as explained in Box 2 of the main text. We always take the electron donor flux as reference, and from the flux $f_i^{\pm}$ one can define the yield $y_i \equiv f_i^{\pm}/f_{\rm ed}^{+}$, analogous to the biomass yield. Even heat fluxes are often reported as heat yields, i.e. using $y_{\dot q} \equiv \dot q/f_{\rm ed}^{+}$, where $\dot q$ is defined in Box 1 of the main text and below in Eq. (B17). Furthermore, with this definition the dissipation, $\sigma$, is nothing but the dissipation yield, i.e. $\sigma = y_{\dot g_{\rm diss}} \equiv -\dot g_{\rm diss}/f_{\rm ed}^{+}$, where $\dot g_{\rm diss}$ is defined in Box 1 Eq. (3). Importantly, this allows to treat in a consistent manner both, chemostat and batch experiments, Fig. SI1. For this reason, in our work we used yields, instead of fluxes, to characterize the cellular growth. See however Extended Data Fig.8 for a description of chemostated data using fluxes.

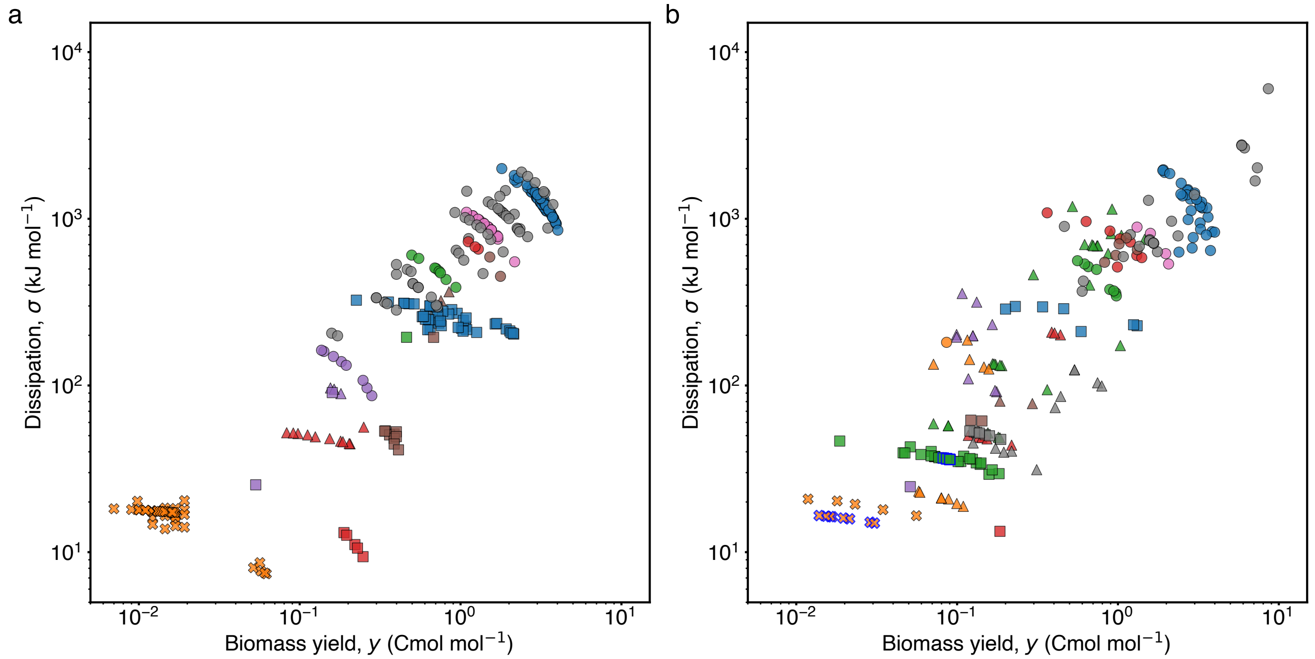


Fig. SI1. **Chemostated and batch cultures exhibit the same behavior. a.** Subset of Fig 3a for chemostated cultures. **b.** Subset of Fig 3a for batch cultures. Blue edges: fed-batch. While our theoretical framework is suited to chemostated cultures and batch cultures arrested in exponential phase, we have also used it on batch cultures allowed to saturate, where it constitutes a rougher approximation. As one can see, both sets of data exhibit similar behavior, showing that our conclusions are robust to difference in growth conditions.

##### 3. Element conservation and macrochemical equation

Each chemical species is characterized by its element composition. The matrix of entries $e_{ik}$ contains this information, as it expresses the number of elements of type $k$ (C, H, O, N, ...) present in chemical species $i$ (e.g. glucose, acetate, ammonia, etc). Since chemical reactions preserve elements, we have that $\sum_i e_{ik} a_{ij} = 0$. Using this in Eq. (B3), we obtain a balance between the element fluxes towards the colony, and those away from the colony (including dilution):

$$\sum_{\mathrm{i} \in \mathrm{sb}} e_{ik} f_i^+ = \sum_{\mathrm{i} \in \mathrm{pd}} e_{ik} f_i^- + \gamma \rho_k \quad , \tag{B15}$$

where the first/second sum runs over substrates/products, and $\rho_k = \sum_i e_{ik} c_i$ refers to the concentration of elements in the colony. Importantly, Eq. (B15) constitutes a set of constraints (as many as elements) on the biomass exchange fluxes, which can be used to determine them Roels [8]. Equation (B15) can be written in terms of yields dividing both sides by $f_{\mathrm{ed}}^+$, which gives Eq. (4) of the main text.

More specifically, each entry in the dataset involves a relatively small number of chemicals exchanged, $C$. This number is typically of the same order and larger than the set of elements spanned by all substrates and products combined, $E$. For example, microbial growth through glucose respiration can be schematically summarized as the following *macrochemical equation*:

$$\mathrm{C_6H_{12}O_6} + \mathrm{O_2} + \mathrm{NH_3} \rightarrow \mathrm{biomass} + \mathrm{CO_2} + \mathrm{H_2O} \quad , \tag{B16}$$

where the above "equation" is not stoichiometrically balanced. In this example, the number of chemicals exchanged is $C = 5$ (3 substrates: glucose, oxygen and ammonia; and 2 products: carbon dioxide and water), and the number of elements is $E = 4$ (C, H, O and N). Eq. (B15) then constitutes a set of four equations, one per element, with six unknowns: the five exchange fluxes and the biomass growth. Provided the measurement of two fluxes, e.g. the biomass growth rate and glucose consumption rate, the rest can be determined by solving the linear system Eq. (B15).

Most of the primary sources that we used to construct the dataset contain very sparse information on the exchange fluxes, and so element conservation was used to complete unknown fluxes in this manner. Furthermore, since the fluxes are often given directly in the form of yields, we solved the linear system for yields rather than fluxes, as discussed in Box 2 of the main text. When redundant information was available, this was used to verify the accuracy of element conservation, see section I.

##### 4. Thermodynamics

Having discussed mass and element conservation, we now construct a non-equilibrium thermodynamic description under steady balanced growth conditions, i.e. conditions under which Eq. (B3) applies. This is the minimal framework introduced in Box 1 of the main text. Assuming knowledge of matter fluxes and thermodynamic properties of biomass and chemicals, the steady state formulation of the second law takes the form:

$$\dot{s}_{\mathrm{prod}} = -\frac{\dot{q}}{T} + \dot{s}_{\mathrm{ch}} + \gamma s \geq 0 \quad , \tag{B17}$$

where $\dot{s}_{\mathrm{prod}}$ is the rate at which entropy is produced, $\dot{q}$ the rate at which heat is produced (or absorbed, if positive), $T$ the temperature set by the media, $\dot{s}_{\mathrm{ch}}$ the rate at which chemical entropy is exchanged, and $\gamma s$ the amount of entropy produced in the form of biomass per unit time, with $s$ the biomass entropy density (see [9] for a comprehensive treatment of thermodynamics of chemical reaction networks). All fluxes are expressed per unit volume of biomass, unless otherwise specified. The rate of chemical entropy exchange is given by

$$\dot{s}_{\mathrm{ch}} = \sum_{i \in \mathrm{pd}} f_i^- s_i - \sum_{i \in \mathrm{sb}} f_i^+ s_i \quad , \tag{B18}$$

and it expresses a balance between the fluxes of entropy due to incoming and outgoing chemicals, with $s_i$ the molar entropy of the corresponding chemical species.

So far Eq. (B17) is incomplete, as it requires an expression for the heat. In other words, it can not be directly used to compute entropy production. To obtain an expression for the heat we make use of Hess' law, which connects heat with enthalpy exchanges. At steady state, this gives

$$\dot{q} = \dot{h}_{\mathrm{ex}} + \gamma h \quad , \tag{B19}$$

where $\gamma h$ is the enthalpy flux due to biomass synthesis and $\dot{h}_{\rm ex}$ is the exchange enthalpy, given by

$$\dot{h}_{\rm ex} = \sum_{i\in{\rm pd}} f_i^- h_i - \sum_{i\in{\rm sb}} f_i^+ h_i \quad , \tag{B20}$$

with $h_i$ the enthalpy of the chemical species. We note that Eq. (B19) is a direct expression of energy conservation, and thus a reformulation of the first law of thermodynamics.

Putting together the expressions of the first and second law, and using the definition of chemical potential, $\mu_i = h_i - Ts_i$, we arrive at

$$T\dot{s}_{\rm prod} = \underbrace{\sum_{i\in{\rm sb}} f_i^+ \mu_i}_{\dot{g}_{\rm sb}} - \underbrace{\sum_{i\in{\rm pd}} f_i^- \mu_i}_{\dot{g}_{\rm pd}} - \gamma g \geq 0 \quad , \tag{B21}$$

which decomposes the total rate of free energy dissipation, $\dot{g}_{\rm diss} = -T\dot{s}_{\rm prod}$, in terms of free energy influx of substrates, $\dot{g}_{\rm sb}$, free energy outflux of products, $\dot{g}_{\rm pd}$, and free energy flux due to biomass synthesis, $\gamma g$, with $g = h - Ts$ the free energy of biomass. In a substantial portion of the dataset growth rates are not reported, and instead only yields are provided. We therefore introduce the dissipation per electron donor, $\sigma = T\dot{s}_{\rm prod}/f_{\rm ed}^+$, which in the main text is simply referred to as dissipation. The second law can then be written in terms of yields as

$$\sigma = \underbrace{\sum_{i\in{\rm sb}} y_i \mu_i}_{-\sigma_{\rm sb}} - \underbrace{\sum_{i\in{\rm pd}} y_i \mu_i}_{-\sigma_{\rm pd}} - yg/\rho_{\rm C} \quad , \tag{B22}$$

where $y_i = f_i^{\pm}/f_{\rm ed}^+$ are the yields of all exchanged chemicals and $y_{\rm ed} = 1$.

Remarkably, Eq. (B19) provides a prediction for growth heat in terms of enthalpy of chemicals and exchange fluxes, including biomass. This prediction can be tested through calorimetry experiments, as is shown in Fig. 1b. We also note that $\dot{s}_{\rm prod}$ and $\dot{q}$ are determined solely by the fluxes of exchanged chemicals, their enthalpies/entropies, the enthalpy/entropy of biomass and the growth rate. Therefore, under steady state conditions, a complete thermodynamic description is possible without any knowledge of the internal reaction fluxes $\nu_j$. We conclude that the thermodynamic properties of cellular growth are determined by three sets of data: the fluxes of matter, which we discussed in B 1 and B 3; the thermodynamic properties of the corresponding chemicals, i.e. their free energy of formation, $\mu_i$, and enthalpy of formation, $h_i$, which we will discuss in H 2; and the thermodynamic properties of biomass, which we will discuss in  H.

##### Appendix C: Defining the cost across and the coefficient within metabolic types

In the main text we grouped the data in terms of metabolic types, $t = 1, \ldots, T$, with each type $t$ containing several instances of data, $d_t = 1, 2, \ldots, D_t$. The metabolic type, also referred to in the literature as macrochemical equation, denotes the set of substrates and products of cellular growth. For a particular type, different data refer to different species, the same species at different temperatures or dilution rates, or experimental variation. In Extended Data Fig.2 we provide a comprehensive list of all metabolic types that the database contains with legends used in figures Extended Data Fig.5, Extended Data Fig.6 and Extended Data Fig.9. In the main text, for simplicity, only some types were highlighted and the index notation was dropped.

With this grouping, the biomass yield of a single data point can be denoted as $y(t, d)$, which refers to the $d^{\text{th}}$ instance of data in the type $t$. Similarly, the dissipation of a data point can be denoted as $\sigma(t, d_t)$. We computed the mean dissipation of a type as $\bar{\sigma}(t) = \sum_{d_t=1}^{D_t} \sigma(t, d_t)/D_t$, and similarly for the mean yield, $\bar{y}(t)$. The yield of a particular data point can therefore be written as $y(t, d_t) = \bar{y}(t) + \delta y(d_t)$. Using these definitions, we identified the across-type cost as $\alpha(t) = \bar{\sigma}(t)/\bar{y}(t)$. Within a type, element conservation establishes a linear relation between $\delta\sigma(d_t)$ and $\delta y(d_t)$, or quasi-linear in case of multiple metabolic products (see B 3). Therefore, we defined the within type coefficient $\beta(t)$ by performing a linear regression of $\delta\sigma(d_t)$ against $\delta y(d_t)$. Our two key findings are simply that $\alpha(t)$ is highly conserved across types, while $\beta(t)$ takes two different values depending on whether the electron acceptor is oxygen or not.

#### Appendix D: Alternative definitions for the thermodynamic cost of growth

In this work, we have defined the cost of cellular growth using a clear non-equilibrium thermodynamic model, see section B. In particular, the cost $\alpha$ is the amount of free energy dissipation per carbon mole of biomass synthesized. This is in the spirit of classical works such as Von Stockar et al. [10] and reference within. We remark that the method used in this work does not introduce putative catabolic and anabolic reactions to compose the macrochemical equation, contrary to what is usually done in the aforementioned body of literature. Our use of the terms catabolic and anabolic follows *after* the computation of the dissipation. Indeed, our definition is based on whether a fundamental force drives or opposes dissipation, and does not refer to intracellular processes. We will now compare our definition and value of cost to alternatives proposed in the literature. To this end, it is useful to recast the value we obtained, $\alpha \approx 500\,\mathrm{kJ\,Cmol^{-1}}$, in different units (see Table I).

| Energy units / per biom. grown | kJ | $k_\mathrm{B}T(= RT/N_\mathrm{A})$ | ATPmol$(= N_\mathrm{A}\mathrm{ATP})$ | ATP (#) |
|---|---|---|---|---|
| carbon mole ($\mathrm{Cmol^{-1}}$) | $500\,\mathrm{kJ\,Cmol^{-1}}$ | $1.25\cdot 10^{26}\,k_\mathrm{B}T\,\mathrm{Cmol^{-1}}$ | $8\,\mathrm{ATPmol\,Cmol^{-1}}$ | $5\cdot 10^{24}\,\mathrm{ATP\,Cmol^{-1}}$ |
| gram ($\mathrm{g^{-1}}$) | $20\,\mathrm{kJ\,g^{-1}}$ | $5\cdot 10^{24}\,k_\mathrm{B}T\,\mathrm{g^{-1}}$ | $0.3\,\mathrm{ATPmol\,g^{-1}}$ | $1.8\cdot 10^{23}\mathrm{ATP\,g^{-1}}$ |
| carbon atom ($\mathrm{C^{-1}}$) | $8\cdot 10^{-22}\,\mathrm{kJ\,C^{-1}}$ | $210\,k_\mathrm{B}T\,\mathrm{C^{-1}}$ | $1.3\cdot 10^{-23}\mathrm{ATPmol\,C^{-1}}$ | $8\,\mathrm{ATP\,C^{-1}}$ |
| cell ($\mathrm{cell^{-1}}$) | $6\cdot 10^{-12}\,\mathrm{kJ\,cell^{-1}}$ | $1.5\cdot 10^{12}\,k_\mathrm{B}T\,\mathrm{cell^{-1}}$ | $1\cdot 10^{-13}\,\mathrm{ATPmol\,cell^{-1}}$ | $6\cdot 10^{10}\,\mathrm{ATP\,cell^{-1}}$ |

TABLE I. **Unit conversion of the dissipative cost of growth ($\alpha$).** Unit conversion of $\alpha \approx 500\,\mathrm{kJ\,Cmol^{-1}}$ to various energy scales: kilo Joules (kJ), thermal energy ($k_\mathrm{B}T$), moles of ATP hydrolyzed (ATPmol), and individual ATP hydrolysis (ATP). Energy scales were normalized either per carbon mole of biomass (Cmol), per gram of biomass (g), per carbon atom of biomass (C), and per cell. The following conversion factors were used: $k_\mathrm{B}T \approx 4\cdot 10^{-21}$ J (and thus $RT \approx 2.45\,\mathrm{kJ\,mol^{-1}}$), hydrolysis of one ATP molecule $\approx 25\,k_\mathrm{B}T$, hydrolysis of one mole of ATP $\approx 60\,\mathrm{kJ}$, average molar biomass weight discussed before $m_\mathrm{b} \approx 24.5\,\mathrm{g\,Cmol^{-1}}$, and Avogadro's constant $N_\mathrm{A} \approx 6\cdot 10^{23}$. Per cell normalization was achieved by assuming an average cell size of $1\,\mu\mathrm{m}^3$ with a dry weight of 0.3 pg, similar to an *E. coli* cell (BNID:103904).

These unit transformations (see Table I) will help us relate the magnitude of our thermodynamic cost of growth to alternative ones introduced in the literature.

1. **The cost as dissipation in the metabolic network.** Whereas early attempts at modeling metabolism using Flux Balance Analysis are thermodynamically inconsistent Varma and Palsson [6], the solution to enforce the preservation of the second law was soon noted beard2002energy. Using this approach, Niebel et al. [1] studied the free energy dissipated by the whole metabolic network of aerobically fermenting *S. cerevisae* and *E. coli.* The authors arrived at maximal values of $\approx 9\,\mathrm{kJ\,g^{-1}}$ for *S.cerevisae* and $\approx 12\,\mathrm{kJ\,g^{-1}}$ for *E.coli.* In our database, for analogous conditions and microbial species, we find values ranging from the same, up to three or four folds higher. Using Flux Balance Analysis on a model of *E. coli* growing on glucose minimal media, Mori et al. [11] computed the flux of ATP production in aerobic and anaerobic conditions. These fluxes correspond to $\approx 7$ and 3 kJ per gram of biomass, respectively. In our database, for the same species in analogous conditions and with similar biomass yield, we find values of dissipation per biomass $\approx 16$ and $7\,\mathrm{kJ\,g^{-1}}$, respectively.

2. **The cost of growth as ATP equivalents.** The main cellular energy currency is adenosine-triphosphate (ATP). Under physiological conditions, ATP hydrolysis to adenosine diphosphate (ADP) and inorganic phosphate results in $\approx 50-60\,\mathrm{kJ\,mol^{-1}}$ of free energy (BNID: 101964). Many metabolic reactions and pathways that transform energy and mass for cellular growth have been biochemically characterized in the last decades. This knowledge allows for experimental and theoretical estimation of the cost of growth in terms of ATP molecules being hydrolized. However, some biochemical reactions do not use ATP as a main energy source, e.g. protein translation uses guanine-triphosphate (GTP) instead of ATP. Similarly, cellular oxidation and reducing power required for growth is provided in the form of adenine-dinucleotides such as NAD(H), NADP(H), and FAD($H_2$). For simplicity, ATP costs are usually estimated as ATP equivalents by converting other currencies such as GTP and adenine dinucleotides to ATP equivalents.

   There are two complementary approaches to estimate the ATP growth requirements – one theoretical and one experimental. The theoretical approach uses biochemical knowledge of metabolic pathways to calculate the amount of ATP equivalents required to build and polymerize precursors to form cellular macromolecules compatible with the known composition of a cell. These theoretical estimates can be compared to experimental methods that commonly use ATP yields ($Y_\mathrm{ATP}$) as a measure of the cost of growth. $Y_\mathrm{ATP}$ is defined as the amount of biomass produced per amount of ATP hydrolyzed and its calculation depends on how much ATP is produced per substrate consumed for a given metabolic type. The notion of $Y_\mathrm{ATP}$ was first introduced byBauchop and Elsden [12], and refined by Stouthamer [13] to consider the cost of maintenancePirt [14].

- **Experimental.** Soon after its definition, an experimental value of $Y_{\mathrm{ATP}} \approx 10.5\,\mathrm{g\,ATPmol^{-1}}$ has been generally accepted Stouthamer and Bettenhaussen [15]. Further experiments and calculations by Stouthamer and Bettenhaussen [15] extended the range of values from $\approx 2$ to $12\,\mathrm{g\,ATPmol^{-1}}$. A comparison with $1/\alpha \approx 3\,\mathrm{g\,ATPmol^{-1}}$ shows that our measure of cost tends to be compatible or higher by a few folds.

- **Theoretical.** Stouthamer [13] provided a theoretical estimate of $Y_{\mathrm{ATP}} \approx 28.8\,\mathrm{g\,ATPmol^{-1}}$, or $\approx 51\,\mathrm{kJ\,Cmol^{-1}}$, starting from glucose and inorganic salts and excluding maintenance. The value changes to $\approx 10.5\,\mathrm{g\,ATPmol^{-1}}$ or $\approx 140\,\mathrm{kJ\,Cmol^{-1}}$ when acetate is the substrate. More recent estimates in the same spirit, e.g. Phillips et al. [16] p.199, have shown the cost of *E. coli* growth under aerobic conditions to be $\sim 10^{10}\,\mathrm{ATP\,cell^{-1}}$ or $\sim 80\,\mathrm{kJ\,Cmol^{-1}}$, excluding maintenance. Although a direct comparison suggests that our result is an order of magnitude larger, we stress that our definition of cost encompasses the overall dissipation of the growth process, maintenance included. tlynch2015bioenergetic use a conversion factor to transform volume into ATP yield. This was done so to partition the cost of a gene, a concept introduced in Lane and Martin [17]. Such conversion factors are common in the case of oxygenic metabolisms, often referred to as P:O ratio. The cost obtained in this manner in Lynch and Marinov [18] for *E. coli*, $\approx 3 \cdot 10^{10}\,\mathrm{ATPcell^{-1}}$ is similar, smaller by a factor of two, to what we obtained. These estimates take into account explicitly the maintenance cost, and implicitly the opportunity cost of cellular metabolism Mahmoudabadi et al. [19].

3. **The cost as free energy required by biomass formation.** In an early attempt, Morowitz [20] estimated the entropy and enthalpy changes of transforming simple substrate molecules into biomass. This gives the value of the free energy required by microbial biosynthesis. Considering a minimal media, Morowitz writes the coarse-grained chemical reactions to form cellular building blocks, such as amino acids and nucleotides, from the substrates available in the media. This allows to estimate the net enthalpy and entropy change of synthesizing the precursors required for 1 g of biomass, on the basis of a representative biomass composition. By further using polymer statistics and known hydrolysis enthalpies of peptides and sugar-phosphate bonds, he estimates the entropy and enthalpy change of those precursor polymerizing into macromolecules. Such estimate, for the anaerobic synthesis of *E. coli* biomass from glucose, results in a free energy change $\approx -1\,\mathrm{kJ\,g^{-1}}$, denoting an exergonig process. More recently, in the spirit of Morowitz and using similar approaches of precursor reaction thermodynamics, McCollom and Amend [21], Amend et al. [22] estimated the energy requirement of aerobic and anaerobic chemolithoautotrophic synthesis of monomeric cellular components. They found the minimal free energy required to amount to $\approx 18$ and $1\,\mathrm{kJ\,g^{-1}}$ of biomass, for aerobic and anaerobic conditions respectively. McCollom and Amend [21] suggest that the inclusion of all anabolic processes would increase the energy requirement and reduce the difference between the aerobic and anaerobic results. Their values of free energy required by anabolism are therefore compatible with our results.

4. **The cost from the macrochemical equation approach.** Several works have used the macrochemical equation approach to study thermodynamic aspects of microbial growth. Although these works and our both rely on the notion of macrochemical equation, the theoretical foundation for this latter is fundamentally different. In most cases, the thermodynamics cost of growth have been implicitly defined as a combination of a catabolic and an anabolic reaction Von Stockar and Liu [23], Battley [24], Kleerebezem and Van Loosdrecht [25], Calabrese et al. [26], Heijnen and Van Dijken [27], Smeaton and Van Cappellen [28], Liu et al. [29], Roden and Jin [30]. There, the catabolic and anabolic reaction are postulated on the basis of biochemical intuition. The net free energy change of biomass growth is then computed balancing one or two Smeaton and Van Cappellen [28], Kleerebezem and Loosdrecht [31] parameters, that account for the ratios of both the anabolic and catabolic reaction. The results of these approaches, which often used a "standard" biomass stoichiometry, suggest a cost of $\approx 300 - 600\,\mathrm{kJ\,Cmol^{-1}}$, in agreement with our estimates, see Fig. 3B in Smeaton and Van Cappellen [28] and Table VG in Heijnen and Van Dijken [27]. We remark that we went to the primary references of these works, and many of these references constitute a subset of our database.

5. **The cost as free energy of formation of biomass.** Another definition of the cost of growth is the free energy of formation of biomass, defined as the energy change of a putative chemical reaction of forming biomass from its constituting elements. Estimating the free energy of formation of biomass is experimentally very challenging, involving measurements of the heat released by biomass combustion to calculate the enthalpy of formation, and low-temperature differential scanning calorimetry to estimate its entropy content. In our work, we incorporated the free energy of biomass formation as part of the cost, and found that its contribution is very small (typically $\leq 10\%$). Indeed, Battley [32] and McCollom and Amend [21] estimated that $\leq 10\%$ of the total free energy is conserved in biomass or cellular protein as a proxy for total biomass. For more detail on the free energy of formation of biomass see section H.

Taken together, the above-mentioned approaches all rely on particular assumptions and mostly underestimate the dissipative cost of growth by a factor of $2-10$. Our approach is based on a formal notion of cost in thermodynamics, i.e. the amount of free energy that is dissipated (or total entropy produced). It does not require any conversion factors based on oxygen respiration or substrate-phosphorylation, nor does it rely on details of the metabolic pathway or arbitrary formulations of anabolic reactions, all details that are often not known. Instead, our method does depend on metabolism through the known substrates and products (see Extended Data Fig.5), their chemical properties (see H 2), and the biomass yield and composition. The content of biomass, both its enthalpic and entropic contributions, is included and species-specificity is considered by its stoichiometric composition, see H. Our approach naturally encompassed different unicellular sizes and doubling times, as the final results are given per unit biomass synthesized. Converting to size and time is straightforward. While some of these features of our model are shared with some of the aforementioned approaches, none of them combine them all.

#### Appendix E: The origin of the cost across types, $\alpha$, and of the within type coefficient, $\beta$

##### 1. Analytical relation between yield and dissipation

The total dissipation in a non-equilibrium system can be reduced to a sum of products between independent fluxes and corresponding generalized thermodynamic forces. For the case of chemical systems, this can be done by identifying conserved quantities , which in our case correspond to elements. We now develop this decomposition.

Following B 3, we label with $i_\mathrm{i}$ the $C-E$ independent yields and $i_\mathrm{d}$ the remaining dependent ones. Since the choice is arbitrary, we chose to treat as independent yields the measured ones, which almost always include biomass. Once the element matrix is restricted to the dependent species only (entries $e_{i_\mathrm{d},k}$), it can be inverted (entries $(e^{-1})_{i_\mathrm{d},k}$). Using element conservation, Eq. (B15), to express dependent yields as a function of independent ones, we arrive at

$$y_{i_\mathrm{d}} = E_{i_\mathrm{d},\mathrm{ed}} - yE_{i_\mathrm{d}} + \sum_{i_\mathrm{i}\in\mathrm{sb}} y_{i_\mathrm{i}} E_{i_\mathrm{d},i_\mathrm{i}} - \sum_{i_\mathrm{i}\in\mathrm{pd}} y_{i_\mathrm{i}} E_{i_\mathrm{d},i_\mathrm{i}} \quad , \tag{E1}$$

where $E_{i_\mathrm{d},i_\mathrm{i}} = \sum_k (e^{-1})_{i_\mathrm{d},k} e_{i_\mathrm{i},k}$, $E_{i_\mathrm{d}} = \sum_k (e^{-1})_{i_\mathrm{d},k} \rho_k/\rho_\mathrm{C}$, and we assumed $i_\mathrm{d} \in$ pd (in the case $i_\mathrm{d} \in$ sb signs are opposite). Using Eq. (E1) in Eq. (B22) we obtain an expression for the dissipation as a function of the independent yields (Eq. (12) of the main text),

$$\sigma = r_\mathrm{ed} - yr_\mathrm{b} + \Delta \quad ,\text{with} \quad \Delta = \sum_{i_\mathrm{i}\in\mathrm{sb}} y_{i_\mathrm{i}} r_{i_\mathrm{i}} - \sum_{i_\mathrm{i}\in\mathrm{pd}} y_{i_\mathrm{i}} r_{i_\mathrm{i}} \quad . \tag{E2}$$

The coefficients $r_{i_\mathrm{i}} = \mu_{i_\mathrm{i}} - \sum_{i_\mathrm{d}} E_{i_\mathrm{d},i_\mathrm{i}} \mu_{i_\mathrm{d}}$, with dimensions of a chemical potential, relate the independent yields $y_{i_\mathrm{i}}$ to the dissipation (there are analogous expressions to those of $r_{i_\mathrm{i}}$ for the biomass coefficient, $r_\mathrm{b}$, and the electron donor coefficient, $r_\mathrm{ed}$). From a thermodynamic point of view, Eq. (E2) is an expression of the dissipation as a product of generalized and independent fluxes (the yields) and forces (the coefficients) Wachtel et al. [33], De Groot and Mazur [34]. We note that for the cases in which there is a single independent yield we have $\Delta = 0$, and so

$$\sigma = r_\mathrm{ed} - yr_\mathrm{b} \quad . \tag{E3}$$

Next, we will relate these expressions of the dissipation to the dissipative cost $\alpha$ and to the coefficient $\beta$ used in the main text.

##### 2. Variations of yield and dissipation across and within metabolic type

To gain further insight into how the decomposition above relates to variation within and across metabolic types, it is important to remark the different sources of data variation. Across types, variations are due to different metabolic pathways being activated or deactivated in presence of different nutrient sources. In contrast, variations of yield for a given type arise from diverse sources: changes in chemical concentrations (that affect the chemical potentials, although weakly, see SI G), changes in temperature (that also affect the chemical potentials weakly through Eq. (G2), with a few percent effect), changes in microbial species using the same metabolic type (which affect the biomass composition and through it the stoichiometry of the equation and the free energy of biomass, also with minor consequences, see Fig. SI2), or experimental variability. Therefore, the sources of variations within and across types are very different, and have different effects on the parameters and variables in Eq. E2.

Taking the above into account, we now decompose within and across variability for the simple case of a single independent yield. In this case, the mean dissipation of a type $t$ is given by

$$\bar{\sigma}(t) \approx r_\mathrm{ed}(t) - \bar{y}(t) r_\mathrm{b}(t) \quad , \tag{E4}$$

where the approximation arises from $r(t,i) \approx r(t)$, and thus we are ignoring changes in temperature and biomass composition within type, which as we said are small. Using Eq. E4 we then have that variations in dissipation within type are given by

$$\delta\sigma(i_t) \approx -r_\mathrm{b}(t)\delta y(i_t) \quad . \tag{E5}$$

This second expression explains the origin of the linear relationships for yield-dissipation variations within metabolic type observed in Fig.s 3c and Fig. 5, and displayed in Extended Data Fig.9. Overall, the linear relationship within type arises directly from element conservation for the case of a single independent yield, due to the small variability on driving forces. In the case of multiple independent yields, deviations from this linearity are expected and observed, and these deviations reflect the role of the additional yields, which is often also small.

Taking into account the approximations above, it is now easy to identify the origins of the deviation coefficient $\beta$ and the across cost $\alpha$. For the coefficient, we simply have that $\beta \approx -r_{\mathrm{b}}$, which we emphasize again only depends on stoichiometry and chemical potentials. We note that, in all metabolic types of the database, the force $r_{\mathrm{b}}$ is positive. Therefore, the yield of biomass is a thermodynamically unfavorable *anabolic* flux that needs to be driven by the remaining *catabolic* fluxes. The across cost instead stems from Eq. E4. Since in a large part of the data the second term on the right hand side is sub-dominant, the force $r_{\mathrm{ed}}$ approximates the driving *catabolic* force of growth. We then have that the conservation of the cost across metabolic types arises from the relationship $r_{\mathrm{ed}}(t) \approx \alpha(t)\bar{y}(t)$, which is shown in Fig. 6.

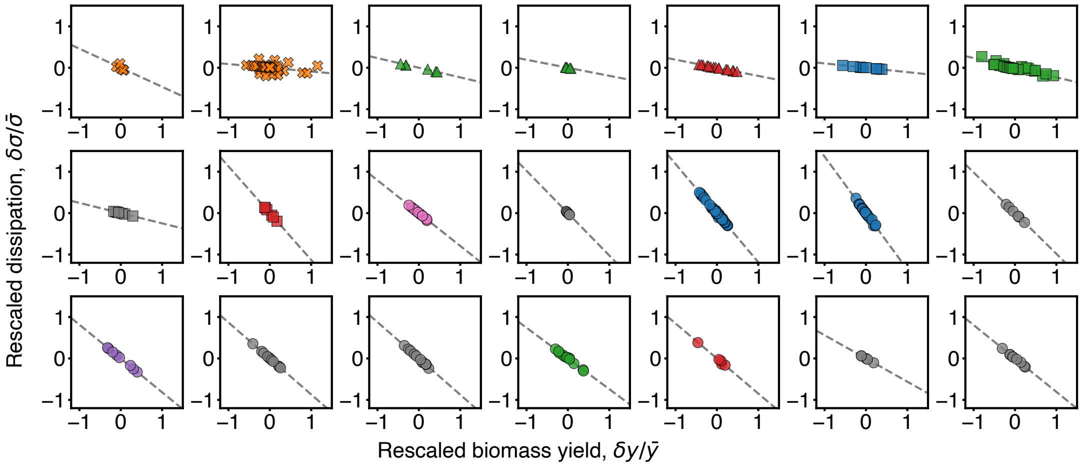


Fig. SI2. **Effect of temperature and biomass composition within metabolic type.** The panels show data for macrochemical equations with one independent yield and more than four data-points per equation. The dashed lines represent the linear relation $y(x) = -\frac{r_{\mathrm{b}}\bar{y}}{\bar{\sigma}}x$, relating the rescaled yield to the rescaled dissipation, see Eq. E5. In each panel, deviations of the data points from the line are due to: changes in temperature affecting the chemical potentials, changes in pH affecting the chemical potential of hydrogen ions, changes in biomass stoichiometry affecting $r$ (through both the free energy of biomass and the element balance of the macrochemical equation). These changes affect both $r_{\mathrm{ed}}$ and $r_{\mathrm{b}}$, and, as one can see, the effect is minor. On the contrary, variations along the line are due to the influence of these parameters on the physiology of the organisms (i.e. on the biomass yield).

##### 3. Efficiency

Having performed the decomposition in Eq. E2, it is possible to define a thermodynamic efficiency for the overall growth process. We follow the approach of Wachtel et al. [33] to define the efficiency as the ratio of the output dissipation over the input dissipation in the following sense. The output dissipation is simply the dissipation in which the generalized thermodynamic forces oppose the corresponding fluxes, and the input dissipation is the opposite. For substrates, input dissipation is the sum of terms with positive force, $r^+_{i_\mathrm{i}} \equiv r_{i_\mathrm{i}} > 0$ for $i_\mathrm{i} \in \mathrm{sb}$, and output dissipation with negative force, $r^-_{i_\mathrm{i}} \equiv r_{i_\mathrm{i}} < 0$ for $i_\mathrm{i} \in \mathrm{sb}$. For products and biomass, the signs are opposite because of the minus sign in Eq. (E2).

In our case, we empirically found that the electron donor force is always positive, $r_\mathrm{ed} > 0$, and the biomass flux opposes its corresponding force, i.e. $r_\mathrm{b} > 0$ (except in three cases). Therefore, we define the efficiency as

$$\eta = -\frac{-y r_\mathrm{b} + \sigma_\mathrm{out}}{r_\mathrm{ed} + \sigma_\mathrm{in}} \quad , \tag{E6}$$

with $\sigma_\mathrm{in} = \sum_{i\in\mathrm{sb}} r^+_{i_i} y_{i_i} - \sum_{i\in\mathrm{pd}} r^-_{i_i} y_{i_i}$ and $\sigma_\mathrm{out} = \sum_{i\in\mathrm{sb}} r^-_{i_i} y_{i_i} - \sum_{i\in\mathrm{pd}} r^+_{i_i} y_{i_i}$. According to this definition, the thermodynamic efficiency is always positive and smaller than one, and it recapitulates the intuition of a coupling between anabolism (in the numerator) and catabolism (in the denominator). Notice that, for the three exceptional cases with negative $r_\mathrm{b}$, the corresponding term appears in the denominator of Eq. (E6).

##### 4. Origin of the differences in deviation coefficient $\beta$ for aerobic and anaerobic types

We now discuss the typical values of $r_\mathrm{b}$, which approximates the coefficient $\beta$, for different metabolic types. For most types, we have that $r_\mathrm{b} \sim 100$ kJ Cmol$^{-1}$. As pointed out in the main text, aerobic respiration has an exceptionally higher value, $\approx 500$ kJ Cmol$^{-1}$. Such a thermodynamic force opposing biomass synthesis implies that a greater biomass yield is achieved with a sharp decrease in dissipation. However, the reduction of oxygen to water is not the only metabolic process associated with a higher value of the force $r_\mathrm{b}$. The metabolic reduction of nitrate, $NO_3^-$, or nitrite, $NO_2^-$, to ammonium, $NH_4^+$, can give $r_\mathrm{b} \approx 300$ kJ Cmol$^{-1}$. Moreover, when the reduction of nitrate to nitrogen, $N_2$, is coupled to the autotrophic reduction of $CO_2$, we find that $r_\mathrm{b} \approx 500$ kJ Cmol$^{-1}$, as in aerobic respiration.

To understand the origin of the greater force opposing aerobic biosynthesis, we can look at the definition given in SI E 1,

$$r_\mathrm{b} = g/\rho_\mathrm{C} - \sum_{i_\mathrm{d}} E_{i_\mathrm{d}} \mu_{i_\mathrm{d}} \quad . \tag{E7}$$

The value of $r_\mathrm{b}$ is the result of a summation. We find the main contribution to come from the chemical potential of carbon dioxide (i.e. bicarbonate), at $\approx -400\,\mathrm{kJ\,mol^{-1}}$ ($\approx -600\,\mathrm{kJ\,mol^{-1}}$ respectively). This role of carbon dioxide extends to the anaerobic respiration cases where $r_\mathrm{b}$ has the exceptionally low values cited above.

However, while necessary, the presence of carbon dioxide is not sufficient to determine the value of $r_\mathrm{b}$. Indeed, carbon dioxide is produced in many anaerobic processes and yet the corresponding value of $r_\mathrm{b}$ is modest. The reason is that each chemical potential $\mu_{i_d}$ in Eq. (E7) is multiplied by the stoichiometric pre-factor $E_{i_d}$. In respiration with oxygen or nitrate/nitrite, $E_{CO_2} = 1$. In most cases of anaerobic fermentation, $E_{CO_2} \ll 1$, suppressing the contribution of the chemical potential. For example, the anaerobic fermentation of glucose to acetate produces carbon dioxide. The pre-factor suppressing the chemical potential of this latter reaches $E_{CO_2} \sim 0.01$ mol Cmol$^{-1}$. Finally, in most cases of anaerobic respiration $E_{CO_2} = 1$, but the effect of carbon dioxide is counterbalanced by the other terms in the sum. For example, when sulfate, $SO_4^{2-}$, is reduced to sulfide, $H_2S$, the contribution of carbon dioxide's free energy is fully counterbalanced by the term due to sulfate itself, with $\mu_{SO_4^{2-}} E_{SO_4^{2-}} \sim 400$ kJ Cmol$^{-1}$.

#### Appendix F: A thermodynamic database of microbial growth

##### 1. Data curation and validation from primary literature sources

Our thermodynamic analysis of growth is based on data compiled from diverse sources. The database contained in the file `thermo_growth_dat.csv` consists of unicellular growth parameters across a wide range of species, environmental conditions, and metabolic types. All data used in this work was collected from published literature, and with minimal exceptions, we used primary sources (see references below). These primary sources were found by extensive literature search, using also as basis previously published meta-analyses such as Smeaton and Van Cappellen [28], Battley [35]. We note that such previous meta-analysis were only used as a source of references, and in all cases the data was curated directly from the original publications. For each experiment, we curated the following information and parameters when available:

- **Organism.** Species name.
- **Growth condition.** Whether the experiments were performed under conditions of continuous growth (i.e. chemostat), fed-batch culture, or batch cultures, see section B 1.
- **Environmental condition.** Chemical constituents involved in growth: carbon source, electron donor, electron acceptor, nitrogen source, and diverse metabolic products (these were indexed $i = 1, 2, \ldots$ in B 1). Also the growth temperature $T$ and pH.
- **Macrochemical equation**. Macrochemical equations characterizing unicellular growth were curated when reported by the literature source, see section B 3. Missing macrochemical equations were constructed using curated information on trophism, metabolic substrates, products, and biomass stoichiometry. Whenever necessary we wrote the corresponding ionization states of chemicals, adding hydrogen ions to conserve charge.
- **Growth rate.** For batch experiments, maximal growth rates during exponential growth, $\gamma$; and for continuous culture experiments, dilution rates were curated, $d$ (with $d = \gamma$, see B 2).
- **Yields and concentrations**. The reported yields of chemical species, $y_i$, including biomass, $y$. In addition, for chemostat experiments, concentrations of biomass or substrate chemicals (together with the dilution rate this allows to compute the chemical fluxes, $f_i^{\pm}$).
- **Stoichiometry**. Stoichiometric element composition of biomass, $e_{\mathrm{b},k}$, see section B 3. When stoichiometric information was available in the primary source, it was parsed. When it was not available, the stoichiometry of the particular unicellular species was taken from a stoichiometry dataset constructed from diverse sources, see H 1. When the species was not available in the dataset, stoichiometry was assumed to be similar to the closest phylogenetic relative (by genus or domain) with available information on its stoichiometry, and matched accordingly.
- **Element recovery**. Whenever element recovery experiments were performed, we annotated the fraction of recovered elements (typically carbon, see Fig. 1a inset and Extended Data Fig.1a inset).
- **Heat**. Calorimetric measurements of heat production, $\dot{q}$, often in the form of a yield $y_{\dot{q}} \equiv \dot{q}/f_{\mathrm{ed}}^{+}$.

As we attempted to find thermodynamic patterns persistent across varied types of experiments, we excluded only a very small fraction of the data that we encountered. We only excluded data when the growth medium was such that we were not able to write down a sensible macrochemical equation for the growth process. We also excluded data when, upon balancing fluxes, there was a substantial violation of the macrochemical equation, with fluxes of particular chemicals taking the opposite sign to that reported in the literature, i.e. products becoming substrates. This for example occurred with data from Hempfling and Mainzer [36], in the case of anaerobic fermentation.

Overall, the database contains 504 data points. Approximately half, 296 data points correspond to continuous culture experiments. The remaining instances correspond to batch cultures with 13 points corresponding to fed-batch culture. 98 data points contained calorimetric heat dissipation measurements, 56 instances reported one or more redundant yields and 51 data on carbon recovery. On occasion, data curation involved digitizing plots, adjusting mistyped units, or the inference of underlying macrochemical equations of unicellular growth. In section F 3, we provide details and comments on the data parsing of each primary literature source. Data curation was extensive and done with great care in order to standardize and convert the different reported units such, that they are directly comparable and usable for the minimal non-equilibrium thermodynamic framework of this work. The units and typical values of variables used in this work are described in the section below, section F 2.

##### 2. Units and typical values of variables used

Multiple choices of units for the thermodynamics of unicellular growth are used in the literature, depending on the particular scientific community. We now make some remarks on the choices of units and provide typical values of the different quantities that we used in this work. Unless explicitly stated otherwise, densities are provided per unit volume of the chemostat, and not per unit volume of the cellular colony. Whereas the latter was used to develop the formalism, both are equivalent at the steady state (results in the main text are given as ratios of densities, so this choice becomes irrelevant).

- **Chemical fluxes**. In chemostat experiments the exchange fluxes are measured from concentrations in the fermentation vessel, i.e. the environment, times the dilution rate. A typical value is $F^+_{\rm ed} \sim 10^{-1}\,{\rm g\,L^{-1}\,h^{-1}}$ for the glucose exchange flux.

- **Biomass stoichiometry**. Biomass is often characterized by a stoichiometric formula, $\mathrm{CH}_{e_{\rm b,H}}\mathrm{O}_{e_{\rm b,O}}\mathrm{N}_{e_{\rm b,N}}\ldots$. Here $e_{{\rm b},k} = \rho_k/\rho_{\rm C}$ measures how many atoms of the corresponding element are present per carbon atom, with $\rho_k$ the elementary concentration defined below Eq. (B15). Typical values for stoichiometry used in the database are $e_{\rm b,H} = 1.7$, $e_{\rm b,O} = 0.5$, $e_{\rm b,N} = 0.2$.

- **"Molar" biomass weight**. It is common to measure biomass in units of carbon moles (Cmol). Provided a stoichiometric formula and the atomic masses, $m_k$, the mass per Cmol of biomass is given by $m_{\rm b} = (m_{\rm C} + m_{\rm H}e_{\rm b,H} + m_{\rm O}e_{\rm b,O} + m_{\rm N}e_{\rm b,N}\ldots)$ with a typical value of $m_{\rm b} = 24.5\,{\rm g\,Cmol^{-1}}$.

- **Biomass**. The biomass density $\rho$ is the amount of dry biomass per volume and is measured in ${\rm g\,L^{-1}}$, whereas the carbon density $\rho_{\rm C}$ is measured in units of ${\rm Cmol\,L^{-1}}$. The former units are sometimes indicated by "g.d.w." (gram dry weight). The biomass concentration in the vessel can be measured sampling a given volume of the media and weighing the corresponding amount of dry biomass. A typical biomass density for a continuous culture of *E. coli* is $\sim 0.1\,{\rm g\,L^{-1}}$ or $\sim 10^{-2}\,{\rm Cmol\,L^{-1}}$, with a vessel volume of $\sim 0.1\,{\rm L}$.

- **Growth rate**. The growth rate $\gamma$ takes values ranging from $\sim 1\,{\rm h^{-1}}$ in fast growing aerobic respiration experiments, to $\sim 10^{-2}\,{\rm h^{-1}}$ for slowly growing cells using inorganic electron acceptors.

- **Yield**. Yields are defined as the ratio of fluxes and their units are dimensionless. Since the fluxes change units in different ways, it does make a difference whether fluxes were measured in grams, moles, or else. To keep the same notation $y$ without ambiguity, the units of the fluxes are often specified as ${\rm g\,g^{-1}}$, ${\rm mol\,mol^{-1}}$, ${\rm Cmol\,mol^{-1}}$, etc.

  For example, the biomass yield is $y = \gamma\rho_{\rm C}/f^+_{\rm ed}$ in Cmol/mol and $Y = \gamma M/\mathcal{M}^+_{\rm ed}$ in g ${\rm g^{-1}}$. The two definitions are related by the change of units $m_{\rm b}/m_{\rm ed}$, see also section B 2. For the case of *E. coli* growing on glucose, we have $y \sim 1\,{\rm Cmol\,mol^{-1}} \sim 0.1\,{\rm g\,g^{-1}}$. In Extended Data Fig.7 we show how changes in the units of yield have little effect in our results.

- **Free energy, enthalpy, entropy**. Thermodynamic potentials are measured in standard units per mol (or per Cmol, in the case of biomass) using free energies, enthalpies, and entropies of formation. For example, the biomass enthalpy of formation is $h \sim -10^2\,{\rm kJ\,Cmol^{-1}}$, as discussed in detail in section H, and the free energy of formation of glucose is $\mu_{\rm gl} \sim -10^2\,{\rm kJ\,mol^{-1}}$.

- **Heat**. The heat density flux, $\dot{q}$, has units of energy per unit volume and time, i.e. power density. In experiments, it is reported the rate of heat produced per biomass grown, with typical values of $\dot{q}/\gamma\rho_{\rm C} \sim 10^2\,{\rm kJ\,Cmol^{-1}} \sim 10\,{\rm kJ\,g^{-1}}$ for *E. coli* growing aerobically on glucose, where the grams refer to dry weight. Because the yield of *E. coli* on glucose is $y \sim 1\,{\rm Cmol\,mol^{-1}}$, we find that $\dot{q}/f^+_{\rm ed} \sim 10^2\,{\rm kJ\,mol^{-1}}$, which is the normalization used in the text to refer to heat as a part of the dissipation, $\sigma$.

- **Cost**. The dissipative cost of biomass of each metabolic type, $\alpha = \bar{\sigma}/\bar{y}$, is measured in units of energy per amount of biomass synthesized, measured in either grams or moles of carbon. A typical value for the cost is $\alpha \sim 500\,{\rm kJ\,Cmol^{-1}}$, which can also be expressed as $\alpha \sim 210\,k_{\rm B}T/{\rm C}$ (per atom of carbon in the biomass). See section D for different unit choices.

##### 3. Comments on data curation and parsing from primary literature

We now provide some comments on the parsing procedure. Due to the diversity of experimental methods and conventions used in the literature, a significant effort was dedicated to standarizing the data. The comments below are not meant to be an exhaustive description of the parsing done in each paper, but to provide the necessary cues to address the peculiarities from each paper. For instance, when multiple yields reported could result in ambiguity, we comment which ones were taken; we specify the constituents for which yields were reported, when these were not biomass; we specify which references contained heat measurements, and how these were obtained; we specify some unit conversions; etc.

- Andersen and von Meyenburg [37]: We obtained yields by dividing reported substrate consumption and product formation yield rate by the growth rate.
- Badziong and Thauer [38]: We parsed the yields at the reported pH optima for growth.
- Badziong et al. [39]: We curated the yields obtained by C-14 acetate labeling (Table 4). Biomass yields were taken from Table 3 reporting the incorporation of $CO_2$ and acetate carbon into biomass.
- Brandis and Thauer [40]: Two experiments were not performed in minimal media and required low concentrations of yeast extract. However, C-14 labelling revealed those sources are not used for energy or as carbon sources and that acetate is fully incorporated into biomass. Biomass yield were reported as intervals. We computed the mean biomass yields (e.g. $9 - 10\,\mathrm{g\,mol^{-1}}$ as $9.5\,\mathrm{g\,mol^{-1}}$).
- Chua and Robinson [41]: This work contains both batch and chemostat experiments at various pH and dilution rates. We calculated the yields using Pirt's equation and data provided in Table 1 using a dilution rate of $D = 0.08\,\mathrm{h^{-1}}$. Minimal media contained additional cysteine.
- Clarens and Molleta [42]: The paper reported product yields (methane on acetate) and growth yield in response to variations of acetate concentration, temperature, and pH. Data points are averages of triplicate measurements which were parsed. From Tables 2&3 and Figures 4&5, we curated the corresponding growth rates.
- Crabbendam et al. [43]: This study uses chemostat cultures to investigate growth in response to variation in glucose concentration, pH, and dilution rate. Product yields are reported as rates. We obtained yields by dividing the reported yield rates by the dilution rates. The experiments in Table 4 contained mannitol in addition to glucose. We assumed glucose as the main electron donor and attribute the total concentration of mannitol and glucose to glucose for simplicity.
- Dijkhuizen et al. [44]: We parsed the yields reported in Table 1. The authors declare a good agreement between predictions based on element balance and measurements. The authors measured and reported biomass stoichiometry and found it to be independent of the dilution rate.
- Esteve-Núñez et al. [45]: We parsed yields and the steady-state concentration of biomass, electron donor, electron acceptor, fumarate, mannitol, and succinate.
- Goldberg et al. [46]: We parsed the yields reported in Table 1.
- Hernandez and Johnson [47]: We curated the yield of biomass and oxygen reported in Tables 1, 3, and 5 as well as cell concentration. We converted the reported generation time, $t_\mathrm{g}$, to growth rate through $\gamma = \ln 2/t_\mathrm{g}$.
- Heyndrickx et al. [48]: Product yields and carbon recovery data from experiments varying dilution rate, pH, and substrate concentration were parsed from Tables 2, 3, and 5. We excluded the yields containing mixed carbon sources and data that was insufficient to solve the stoichiometry of the marco-chemical equation. The media for the batch cultures used in this work was not minimal and therefore we did not parse the data.
- Huser et al. [49]: We parsed the reported yields of methane produced per acetate consumed.
- Ingvorsen et al. [50]: The parsed yield is the average of the reported batch cultures.
- Laanbroek et al. [51]: The basal media contains yeast extract. The biomass is estimated by carbon measurement. We parsed the biomass yields from Table1. We converted units of grams of carbon per moles of electron donor to carbon moles dividing by carbon molecular weight. We also parsed the yields of acetate produce per ethanol consumed.
- Lin et al. [52]: The media contained peptone and thioglycolate. All product yields were measured and we curated them accordingly.
- Lovitt et al. [53]: The minimal media contains essential amino acids and fatty acids. We parsed all chemostat yield data reported in Table 3 as well as the reported carbon recovery.

- Mayberry et al. [54]: We parsed Table 2, which reports averages from repeated experimental runs for concentrations and yields of biomass and oxygen.
- Myers and Nealson [55]: The defined media contains amino acids. We parsed the yield reported in the text and shown in Figure 3A. The yield was obtained by measurements of cell count and converted by the authors to grams of cells assuming a conversion factor of $2.8 \cdot 10^{-13}$ g cell$^{-1}$.
- Patel [56]: The media contains cysteine. We parsed the biomass and methane yields for two acetate concentrations reported in the text.
- Peters et al. [57]: We parsed data from Table 1 of growth on either lactate or $H_2$ at different dilution rates and temperatures.
- Pfennig and Biebl [58]: The authors measured biomass dry weight by cellular nitrogen measurement. We parsed biomass and reduced electron acceptor yields from Tables 2 & 3 as averages from individual reported replicates.
- Robinson and Tiedje [59]: The media contains cysteine. The yield of proteins on electron donor reported is obtained by fitting an integrated Monod equation for substrate utilization. We used a protein content of biomass of 55%, in analogy with Macy and Lawson [60] to obtain biomass yields. For sulfidogens and methanogens, we parsed the initial concentration of $H_2$ taken from Figures 4 and 5 respectively.
- Roden and Lovley [61]: We only parsed yields of electron acceptor reduced per electron donor consumed of cultures growing in the presence of $MnO_2$, in which yeast extract was not present.
- Rutgers et al. [62]: We parsed yields and dilution rates reported in Table 1 which have been calculated using Pirt equation from experimental data. The biomass is expressed in carbon moles and the stoichiometry has been reported in the paper.
- Sanford et al. [63]: Media contains additional cysteine. Measurements of 16S rRNA gene copy numbers served as a proxy for the number of cells. Transformation to dry weight was done by the authors assuming $3.8 \cdot 10^{-14}$ gr RNA cell$^{-1}$. We parsed data reported in Table 1 for growth of *A. dehalogenans* and the two yields for the *Geobacter* species reported in the text (in cell numbers).
- Sass et al. [64]: We parsed the data reported in Table 2.
- Schauer and Ferry [65]: The minimal growth media contained small amounts of yeast extract and cysteine. We parsed the biomass yield for methane and formate.
- Smith and Mah [66]: The minimal growth media contained small amounts of yeast extract and cysteine. We parsed the yields reported in Table 1. Based on the data provided, we assumed acetate as the sole energy and carbon source.
- Zehnder and Wuhrmann [67]: The media contained additional cysteine. The authors conclude cysteine is an essential sulphur source contributing 0.04 mg per mg of biomass. We assumed the contribution of cysteine to the overall thermodynamic and element balance to be negligible and parsed data reported in Table 3 for "enrichment media" (not containing yeast extract).
- Weimer and Zeikus [68]: We parsed yields of biomass over methane produced. $CO_2$ and $H_2$ were reported to be the sole carbon and energy sources.
- Stieb and Schink [69]: We parsed data reported in Table 2. We derived the yield from reported measurements of product and biomass concentrations.
- Strohm et al. [70]: We parsed data from Tables 1, 2 and 3.
- Szewzyk and Pfennig [71]: We parsed data from Table 1 (batch cultures), and Table 2, (chemostat experiments). The authors report a carbon recovery between 85 to 108 percent for performed experiments.
- Tang et al. [72]: We parsed the biomass yields from Table 1 (for chemostat experiments). The authors used biomass stoichiometry data from previous literature.
- Vasiliadou et al. [73]: The media contains tap water (synthetic wastewater). *Acinetobacter* grows in two phases, with nitrite accumulation and consumption. We parsed the yields from Table 5, which were obtained by fitting of the data with a kinetic model by the authors. We converted grams of nitrate nitrogen to moles using the molecular weight of 76 grams per mole. We converted grams of nitrite nitrogen to moles using the molecular weight of 60 grams per mole.

- Wallrabenstein et al. [74]: We parsed data reported in Table 1 containing growth yields, biomass, and product concentrations as well as the reported carbon recovery.

- Widdel and Pfennig [75]: The biomass is measured directly as dry weight. We parsed the data reported in Table 1 reporting differences in concentrations of biomass, electron donor used and electron acceptor reduced at different stages of growth compared to controls. We computed the yield from a least squares errors linear fit of the reported concentrations.

- Widdel and Pfennig [76]: We parsed the data in Table 2 reporting differences in concentrations of biomass, electron donor used and electron acceptor reduced at different stages of growth compared to controls. We computed the yield from a least squares errors linear fit of the reported concentrations.

- Widdel and Pfennig [77]: We parsed the data in Table 2 reporting differences in concentrations of biomass, electron donor used and electron acceptor reduced at different stages of growth compared to controls and computed the yields from the total concentrations of electron donor used. Table 3 contained concentrations of electron donor used and electron acceptor reduced. We parsed these data taking the ratios as yields.

- Yang and Okos [78]: The media contained small amounts of yeast extract. We parsed biomass yield from Figure 3 and text, which was computed from the biomass formed at the end of the exponential phase with respect to acetate consumption.

- Yoon et al. [79]: The dry weight of biomass was obtained by correlating optical density and gene copy number. We parsed data reported in Table 1.

- De Poorter et al. [80]: Media contained additional cysteine. The continuous cultures were operated at steady-state, whereas the fed-batch experiments had lag-exponential-linear phases of growth. Table 3 reports the maximum yields or the yield in the linear growth phase. In the linear phase, which we parsed, the fluxes have the same time dependency, which make them consistent with our framework, see section B 2.

- de Vries et al. [81]: The media contained a small amount of trypsin digested broth. The dry weight of biomass was measured directly. We parsed yields from Table 1.

- Von Stockar and Liu [23]: This paper reviews previously published results by the authors. We parsed two data points reported in Table 5. The original reference is Duboc [82]. This work contained calorimetric heat measurements.

- Brettel et al. [83]: This work reported the overall yield of biphasic aerobic growth on glucose, with production and subsequent utilization of fermentation products such as ethanol. We parsed biomass yields, reported in grams. We converted them to carbon moles using the average value of $\approx 25\,\mathrm{g\,Cmol^{-1}}$ based on a standard biomass stoichiometry. We also parsed heat measurements done with a flow micro-calorimeter.

- Birou et al. [84]: We parsed the yields from Table IV and one data point from Table VII. When product formation was reported, we computed missing yields using the reported carbon recovery (Table VII). In three instances, we further used the reported measurements of heat dissipation yields (energy recovery, ER). We excluded these data points in our validation of Hess' law. Yields using the energy recovery were computed according to the formula: $f_{\mathrm{p}}^{\mathrm{b}} = \frac{(1-\mathrm{ER})h_{\mathrm{ed}}^{\mathrm{c}}}{h_{\mathrm{p}}^{\mathrm{c}} f_{\mathrm{b}}^{\mathrm{ed}}}$, where ER is the value reported in the original paper, $f_{\mathrm{p}}^{\mathrm{b}}$ is the product yield on biomass and $h_{\mathrm{p}}^{\mathrm{c}}$ and $h_{\mathrm{ed}}^{\mathrm{c}}$ are the enthalpies of combustion of product and electron donor, respectively. Heat measurements were performed with a modified bench-scale calorimeter.

- Dejean et al. [85]: We parsed the growth yields and heat measurements reported in the main text and in Fig. 2 of the reference. The yeast has two exponential growth phases, for which data are reported separately. Therefore, we constructed two macrochemical equations for the two phases. We converted the measured biomass from grams to carbon moles using the molecular weight 24.8 g/Cmol and ash fraction reported in the paper. We obtained the products yields from the enthalpy yields using the combustion enthalpies reported, together with the aid of Fig. 3 of the reference. From the calorimetric-respirometric ratio reported in the reference, we could compute the oxygen yield. We compared this latter to the oxygen yield obtained enforcing element conservation.

- Battley [86]: We parsed the yields, biomass composition, macrochemical equation, and heat dissipation measurements corrected for side processes other than growth.

- Dermoun and Belaich [87]: We parsed the reported yields, macro chemical equation, and heat dissipation measurement (performed on a modified differential scanning calorimeter).

- Dermoun and Belaich [88]: We converted the biomass yield, reported in grams, to carbon moles using the reported biomass stoichiometries. We excluded data for growth on acetate due to mismatching enthalpy recovery and undetected product formation reported by the authors. Heat measurements performed by using a modified differential scanning calorimeter were also parsed.

- Belaich and Belaich [89]: We used only data for growth on glucose as other experiments provide no information on product formation. Part of the reported data can be originally found in an accompanying paper Belaich and Belaich [90]. Calorimetry experiments were performed using a modified differential scanning calorimeter. The initial concentrations of substrates are provided as an upper bound. We converted the biomass yield to carbon moles using $22.8\,\mathrm{g\,Cmol^{-1}}$.

- Ishikawa et al. [91]: The yields of biomass and heat reported correspond to ratios of integrated quantities. Of note, the authors ignore two growth phases, first acetate formation and then acetate utilization. The heat measurements were performed with a twin-type heat conduction micro-calorimeter. We remark that using the reported measured heat dissipation in this work to validate the predicted enthalpy dissipation based on the reported yield showed a large deviation. This work and other works by the authors Ishikawa and Shoda [2] have been criticized in Battley [24] for being a scenario of non-balanced growth. Marison and Von Stockar [92] argue that the observed enthalpy balance mismatched are due to undetected metabolic products.

- Ishikawa and Shoda [2]: We digitized Fig. 2 and 4 and parsed Table I of the paper to extract the data. We matched the heat measurements with the biomass/nutrient concentrations according to the dilution rate. We established a threshold of $0.003\,\mathrm{h^{-1}}$ discrepancy in dilution rate to reject a match. At a high dilution rate, the production of acetate is reported but no measurements are provided. Taken together, we only considered two out of 13 reported data points. We converted grams of biomass to carbon moles using Table I of the paper.

- Tamiya [93]: Data was parsed from Battley [35], as there was no access to the primary source. Of note, Tamiya et al. are the first to elaborate on the concept of degree of reduction and anabolic growth equations. The element analysis of biomass was performed by Yamagata [94].

- Whelton and Doudoroff [95]: Data was parsed indirectly from Battley [35].

- Samejima and Myers [96]: Data was parsed indirectly from Battley [35].

- Hoover and Allison [97]: Data was parsed indirectly from Battley [35].

- Liu et al. [98]: We parsed yields and heat dissipation measurements performed with a bench-scale calorimeter. Acetate (electron donor) concentration was maintained constant to a value reported as $5-10\,\mathrm{g\,L^{-1}}$, pulsed in at a very low rate (we used $5\,\mathrm{g\,L^{-1}}$).

- Birou and Von Stockar [3]: We digitized Fig. 1 to obtain concentrations of biomass, heat dissipated (measured with a bench-scale heat flux calorimeter), and residual glucose. We converted the biomass from grams to carbon moles using the molecular weight of *K. marxianus* biomass stoichiometry. We excluded data for the highest dilution rate data due to detection of uncharacterized products.

- Marison and Von Stockar [92]: We parsed the biomass and product yields from Table IV, and heat and concentration of chemicasls from Table II. Heat measurements were performed with a bench-scale heat flux calorimeter. We excluded the first two data points because the calorimeter operated close to the sensitivity limit. We used the biomass composition reported in the paper.

- Patino et al. [99]: We parsed reported yields, heat dissipation, and macrochemical equations with biomass composition. Heat measurements were performed with a modified commercial reaction calorimeter. We excluded data where biomass growth measurements were out of the sensitivity range of the measuring device.

- Zhao et al. [100]: We parsed yields and fluxes of biomass, $CO_2$, and acetate from continuous cultures. The organisms used in the experiments include genetic variants and mutants of the same species.

- Hempfling and Mainzer [36]: We used $24\,\mathrm{g\,Cmol^{-1}}$ to convert the biomass measurements. We took aerobic data from Tables 2 and 3 (converting grams of oxygen to moles). We excluded fermentative data from Table 1 because we could not write a reliable macrochemical equation. Either element conservation was violated, or the second law was violated, or we did not have enough yields measured to determine the system.

- Mainzer and Hempfling [101]: We treated the data analogously to that in Hempfling and Mainzer [36]. We parsed anaerobic data from Table 1 and aerobic data from Table 3.

- Verduyn [102]: We parsed the data on *C. utilis* growth and biomass composition reported in Table 8 p.143. The composition of biomass for *S. cerevisiae* is taken from Eq. 3 in Verduyn et al. [103]. In experiments with alanine and glutamate, we included ammonia as a product to allow for nitrogen balance.
- Bernacchi et al. [104]: Growth media contained additional cysteine, which was described as not being a carbon source. We parsed the balance of carbon, nitrogen, and degree of reduction from Table 1 and used a macrochemical equation based on Schill et al. [5].
- Liu et al. [4]: We parsed reported data by digitizing Fig.s 3, 4A, 4B, 6 and 7. We computed yields from fluxes and took the biomass composition and macrochemical equations from Schill et al. [5].
- Schill et al. [5]: We parsed reported data by digitizing Fig. 1. The heat measurements were performed with a bench-scale heat flux calorimeter.

#### Appendix G: Thermodynamic properties of substrates and products

According to the framework in B 4, a complete thermodynamic description for a growing cellular colony requires detailed knowledge of the free energies of formation for all substrates and products of metabolism (as well as of biomass, see H). The free energy per mole for an aqueous solution of a chemical species $i$ is given by its chemical potential

$$\mu_i = \mu_i^{\mathrm{o}} + RT \ln\left(\gamma_i \frac{c_i}{c^{\mathrm{o}}}\right) \quad , \tag{G1}$$

where $\mu_i^{\mathrm{o}}$ is the standard chemical potential measured in some standard state, $c_i$ is the concentration, $c^{\mathrm{o}}$ the concentration at the standard state, and $\gamma_i$ is the activity coefficient that measures deviations from the ideal approximation ($\gamma_i = 1$ in the ideal case).

The experiments that we study were normally performed under constant pH conditions. Moreover, many chemical species existed in one or multiple ionization states. This has two implications for the precise computation of the chemical potential. First, the activity coefficient $\gamma_i$ deviates from unity due to the presence of Coulomb interactions, and is given by the Debye-Hückel equation for ionic solutions [105]. Second, the existence of multiple ionization states for chemicals requires that either (*i*) the Alberty's method is used for correcting the chemical potential Silbey et al. [105], Alberty [106], Beber et al. [107], or (*ii*) ionized equations are written for each ionization state subspecies keeping track of charge balance and including the pH value in the free energy of the charged hydrogens Cannon and Raff [108], Sabatini et al. [109]. Concerning the Debye-Hückel correction to the free energy, in practice its role is minor. Concerning the role of multiple ions, it is often the case that one ionization state dominates over the rest, so that keeping track of the dominant ion provides a good approximation.

For completeness, in our work we used two different approaches to deal with the role of ionization. On the first approach (contained in the file `thermo_chem.csv`), which is presented in the main text, we neglected the Debye-Hückel correction and considered only the charge-balanced macrochemical equation for the dominant ion. We therefore did not transform the chemical potentials using Alberty's method, and instead included the free energy of hydrogen ions that compensate the charge imbalance, $\mu_{\mathrm{H}^+} = -RT\mathrm{pH}\ln 10$. On the second approach, we used the Alberty's method and therefore included both types of corrections. The uncorrected values of the free energy, (*i*), were taken from the references indicated in `thermo_chem.csv` file. The corrected values of the free energy, (*ii*), were taken from the eQuilibrator database Noor et al. [110], setting the corresponding value of pH, a typical ionic strength of 0.25 M and pMg value of 3. As we lacked the value of the concentrations of chemicals in most instances, we used $c_i = 1\,\mathrm{mM}$ as commonly done in the literature Battley and Dykhuizen [111]. As one can see in Fig. SI3, the differences between both approaches are only quantitative. Furthermore, an increase/decrease in concentration of three orders of magnitude changes the chemical potential by $\approx \pm 17\mathrm{kJ\,mol}^{-1}$, which is often a negligible contribution compared to the value of the chemical potential itself.

Key to the validation of the dataset, we compared the heat dissipation measurements available with the enthalpy balance according to Eq. (B19), see I. To compute the balance, we collected the enthalpies of formation in aqueous solution of substrates and products of metabolism, as well as of biomass (see H). The enthalpies used are reported in the file `thermo_chem.csv` together with the free energies and the reference to the sources of the data. In the choice of the sources, we prioritized using the same source for as many entries as possible, so to guarantee self-consistency within the database. The enthalpies were also used to correct the use of standard free energies at non-standard temperature, according to the Gibbs-Helmholtz equation

$$\mu_i^0(T) = \mu_i^0(T^0)\frac{T}{T^0} + h_i\left(1 - \frac{T}{T^0}\right) \quad , \tag{G2}$$

where $T^0 = 298$ K and $T$ is the actual temperature of the experiment. The effect of such correction is typically small.

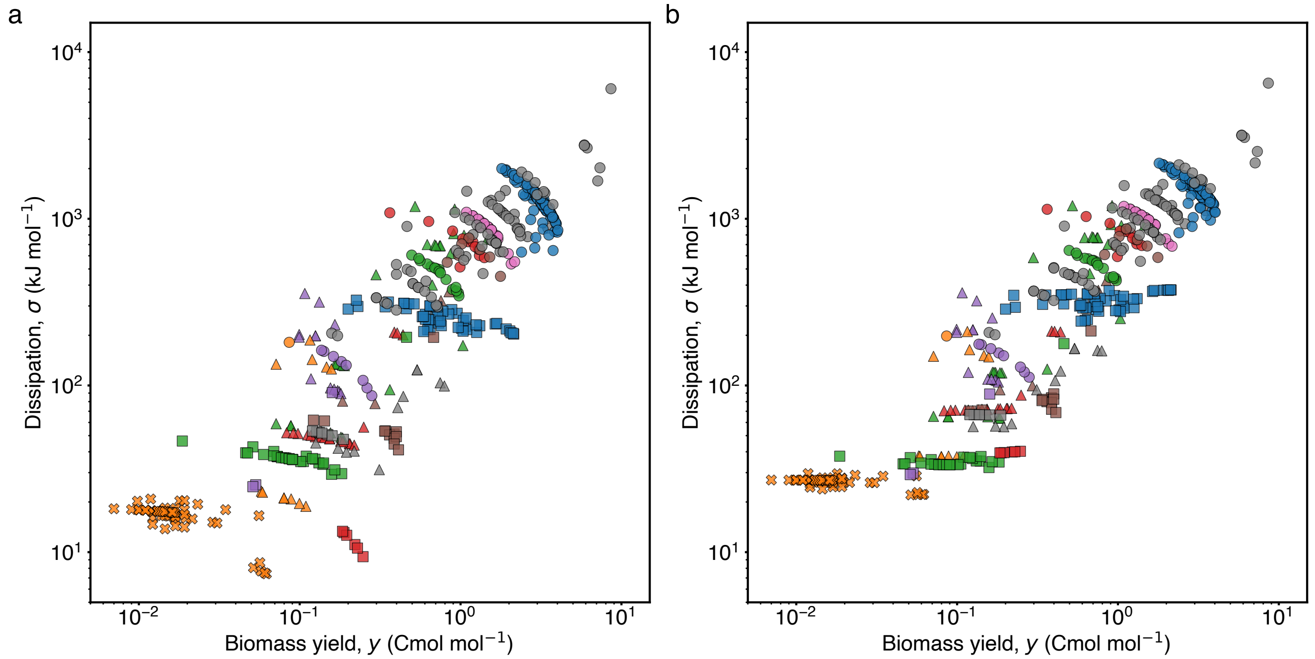


Fig. SI3. **Dissipation against yield using two approaches to correct the chemical potentials for pH changes. a.** Dissipation against yield for the whole dataset, as in Fig. 3a. In this case, the chemical potentials of each chemical was taken from `thermo_chem.csv`, with no correction for the pH of the media. Instead, we balanced charges by using hydrogen ions in the macrochemical equation. The pH correction to the dissipation came by giving a chemical potential to the proton ions related to the pH, as explained in SI G. This was the approach followed in the main text, i.e. this panel is identical to Fig. 3a. **b** Analogous plot to that in a, with a different approach to compute the chemical potentials. In this case, we used the free energies of formation from eQuilibrator Noor et al. [110], which include the pH correction performed using the approach introduced by R.A. Alberty alberty2005thermodynamics. The hydrogen ions were therefore devoid of chemical potential in the dissipation balance. The free energies are taken at a concentration of 1 mM, pMg = 3, ionic strength = 0.25 M and pH of the experiment. When the pH of the experiment was not available in the original reference, we set the value to 7. For bicarbonate, we used the free energy of CO2(total). The data for $MnO_2$, $Mn^{+2}$, S (s) and $C_{16}H_{34}$ were not available in eQuilibrator and therefore we took them from `thermo_chem.csv`. We transformed the free energy of hexadecane according to Alberty's method Alberty [106]. As one can see, both approaches yield similar results.

#### Appendix H: Thermodynamic properties of biomass

##### 1. Biomass composition

The biomass composition is characterized by the concentration of each of its constituents, $c_i$, with $i = 1, \dots, M$, see section B 3. However, as we have seen in section F 2, for our purposes it is more convenient to describe biomass by its elementary composition normalized to carbon content, $e_{\mathrm{b},k} = \rho_k/\rho_\mathrm{C} = \sum_i e_{ik} c_i / \sum_i e_{i\mathrm{C}} c_i$, where $e_{ik}$ is the elementary composition matrix introduced before. The elementary composition of biomass, also referred to as biomass stoichiometry Sterner and Elser [112], has been extensively measured through elementary analysis Gurakan et al. [113]. From these experiments, the elementary composition of different species is available. Because elementary composition of biomass is necessary for our analysis, the data-file `yields.csv` contains a specific biomass composition for each entry. We now describe how these biomass compositions were determined.

For data in which the primary sources described in section F 3 contained specific biomass composition, the corresponding stoichiometry was used, and inserted in our main data-file. However, many of the primary sources we used did not contain biomass stoichiometry data. We therefore compiled a comprehensive dataset of biomass stoichiometry that consists of: (*i*) all the stoichiometric data in the primary sources described in F 3, and (*ii*) the data in the meta-analysis study Popovic [114] that referred to species in our study. Our biomass stoichiometry dataset is in the file `biomass_composition.csv`, and includes dozens of different species. For some of the species there are multiple entries, from measurements performed by different groups or under different conditions. We filled the missing biomass compositions in our main data-file using the following approach: for species in our study that appear in `biomass_composition.csv`, we took the corresponding value (or the average, when multiple values appear in the biomass dataset); and for species in our study that did not appear in `biomass_composition.csv`, we took the average value of the genus or domain in `biomass_composition.csv`. While this approach leads to inaccuracies, the variations in biomass stoichiometry observed across species did not significantly influence the results (Fig. SI2).

##### 2. Enthalpy, entropy and free energy of biomass

The thermodynamic description developed in section B 4 requires knowledge of the enthalpy, $h$, entropy, $s$, and free energy, $g$, of biomass. Because direct measurement of these quantities is challenging, they are often replaced by detailed theoretical estimates Morowitz [20], McCollom and Amend [21], Grosz and Stephanopoulos [115]. Despite this, we now provide an overview of existing measurements for enthalpy and entropy of biomass, and how these can be combined with the elementary composition of biomass to provide empirical estimates for the thermodynamic properties of biomass.

The enthalpy of biomass is obtained by direct measurement of the heat released during the combustion of dry biomass von Stockar et al. [116]. This "heat of combustion", $q_\mathrm{c}$, is also called "enthalpy of combustion", as it refers to the enthalpy difference of the following (unbalanced) combustion reaction:

$$\mathrm{O_2} + \text{biomass} \to \mathrm{CO_2} + \mathrm{H_2O} + \mathrm{N_2} \quad . \tag{H1}$$

Typical values of the heat released are $q_\mathrm{c} \approx 500\,\mathrm{kJ\,Cmol^{-1}}$ for dry biomass, see Table III in von Stockar et al. [116] (as explained in section F 2, for practical purposes biomass properties are better reported on a per Cmol basis.). The heat released during the combustion of organic material has been shown to depend on the stoichiometry of the organic sample, $\{e_{\mathrm{b},k}\}$, and to be well approximated by the following empirical formula:

$$q_\mathrm{c} \approx p\, d(\{e_{\mathrm{b},k}\}) \quad . \tag{H2}$$

Here $d$ measures the number of electrons transferred from the organic material to oxygen, and $p = 115\,\mathrm{kJ\,Cmol^{-1}}$ is an effective proportionality constant Thornton [117]. For biomass, we have that $d = 4e_{\mathrm{b,C}} + e_{\mathrm{b,H}} - 2e_{\mathrm{b,O}}$, which is therefore proportional to the so called degree of reduction, $d/4$, see Cordier et al. [118] and references therein. The validity of this expression, when applied to biomass, has been verified in Cordier et al. [118], Patel and Erickson [119], Gnaiger and Bitterlich [120]. Once the combustion reaction has been determined, the enthalpy of biomass can be obtained by solving the corresponding enthalpy balance:

$$h = a_{\mathrm{N_2}} h_{\mathrm{N_2}} + a_{\mathrm{H_2O}} h_{\mathrm{H_2O}} + a_{\mathrm{CO_2}} h_{\mathrm{CO_2}} - a_{\mathrm{O_2}} h_{\mathrm{O_2}} - q_\mathrm{c} \quad , \tag{H3}$$

with $a_i$ the appropriate stoichiometric coefficients determined by the biomass stoichiometry. Together, Eqs. (H2) and (H3) relate the composition of biomass to its enthalpy. We note that Eq. (H3) requires knowledge of the enthalpy of the chemical species participating in the reaction ($\mathrm{O_2}, \mathrm{CO_2}, \mathrm{H_2O}, \text{and}\, \mathrm{N_2}$). Here, we use the enthalpy of formation for these chemical species, which is the enthalpy change of a putative formation reaction of each molecule. The reactants of these formation reactions, chosen to be the elements in their more stable form at standard conditions, provide a reference (zero) value for the enthalpies Hołyst and Poniewierski [121]. As a result, the enthalpy of biomass will be an enthalpy of biomass formation, with a typical value of $h^{\mathrm{f}} \approx -90$ kJ Cmol$^{-1}$. We also note that we have used measurements of combustion for dry biomass. Clearly, under physiological conditions, biomass is hydrated. However, the enthalpy from hydration is small, measured in von Stockar et al. [116] to be on the order of $q_{\mathrm{c}} \approx 1\,\mathrm{kJ\,Cmol^{-1}}$, and can therefore be neglected.

For the entropy of biomass, direct measurements rely on low-temperature calorimetry of pelleted dry biomass. This determines the heat capacity, which integrated returns the entropy, with a typical value $s \approx 30\,\mathrm{J\,g^{-1}\,K^{-1}} \approx 1\,\mathrm{J\,Cmol^{-1}\,K^{-1}}$ Battley et al. [122]. As hydration entropy has been estimated to have a minor contribution to the overall entropy Battley [123], the value measured for dry biomass, as well as similar ones obtained more recently in the same way Popovic et al. [124], can be used as approximate measurement of biomass entropy. One key point is that these entropy values have been shown to correlate well with the entropy of the elementary constituents of biomass Battley and Dykhuizen [111], Battley [123]. As a result, the following empirical formula that relates biomass entropy to its elementary composition has been proposed:

$$s \approx 0.2 \sum_k \rho_k s_k \quad , \tag{H4}$$

where $s_k$ is the entropy per atom of the corresponding elementary form (e.g., gaseous $H_2$ for hydrogen, graphite for carbon, etc.). In particular, we used $s_{\mathrm{C}} = 5.8$, $s_{\mathrm{H}} = 130.7/2$ , $s_{\mathrm{O}} = 205.1/2$ , $s_{\mathrm{N}} = 191.6/2$, all units of J mol$^{-1}$ K$^{-1}$, taken from Linstrom and Mallard [125]. Because we have used enthalpies of formation, we also will use entropy of formation. We therefore relate the entropy in Eq. (H4) to that of a putative formation reaction. The resulting entropy, which we used in our work, is

$$s^{\mathrm{f}} \approx -0.8 \sum_k \rho_k s_k \quad . \tag{H5}$$

Note that, for consistency, entropies and enthalpies of all chemical species involved in the growth process will also have to be the ones of the formation reaction. To simplify notation, in the main text we omit the superscript. Finally, the free energy of biomass is obtained as $g = h - Ts$, with a typical value of $\approx -40$ kJ Cmol$^{-1}$.

#### Appendix I: Dataset validation

Our work includes a large variety of data that we cross-referenced, including biomass stoichiometric data, growth yield data, and thermodynamic data of chemicals. Furthermore, each of these data types was obtained from a hetergeneous corpus of literature. Due to this diversity of sources and cross-referencing, validation of our dataset is crucial. Three key validation tests were performed: energy conservation, element conservation, and validity of the second law.

- *Energy conservation.* We validated energy conservation using the first law of thermodynamics, which establishes that the heat due to growth equals the balance of enthalpies between substrates and products of metabolism, see Box 1 in the main text and SI B 4. We therefore computed the enthalpy balance for a subset of the data (total of 98 data points) that contained calorimetric measures, and compared it to the reported heat. This comparison was done using per biomass quantities, i.e. yields as described in section B 2. The results of the comparison is shown in Fig. 1b. As one can see, over several orders of magnitude the heat yield predicted through enthalpy balance agrees well with calorimetric measures (see also Extended Data Fig.1 for details on legends). Furthermore, this agreement extends to reported measures of endothermic growth, see inset. We note that in few instances, calorimetry (referred to as "energy recovery") was used to determine specific product yields. These instances, for which there are comments in section F 3, were removed from our validation plot.

- *Element conservation.* In this work we tested element conservation in two different ways. First, we noted the instances in which the number of yields measured (of biomass, products, or substrates) was higher than the number of yields needed to resolve all yields using element conservation (SI B 3). We then used element conservation to predict those yields that had already been measured. The results, which correspond to a total of 69 yields from 56 experiments and 13 different sources, appear in Fig. 1a. As one can see, there is a good agreement, with most deviations occurring for small yields (see also Extended Data Fig.1 for details on legends). Second, we parsed the results of element (carbon) recovery experiments done in some of the growth data that we used (a total of 51 data points). The data appear as a histogram in the inset of Fig. 1a, and show that carbon recovery typically resulted in a good validation of element conservation. As in the case of heat, in few instances the primary sources used element recovery experiments to determine a particular product yield. These instances were removed from our validation plot.

- Finally, while no direct measurements of free energy dissipation have so far been reported in the literature, we tested whether the dissipation $\sigma$ was positive for the collected data. Indeed, we found that 504 out of 509 data points complied with the second law. In 5 cases, the original reference did not report full information about products and substrates of the experiment. The tentative macrochemical equation that we postulated resulted in a negative dissipation and we did not include these data points in our study.

Overall, three different and independent tests were performed to validate the data collected in a total of 302 data points out of the 504 that we used in our study. Given the large variety of data included in this test-set, we find the results of the tests remarkably positive. This validation supports the validity of our approach to the whole dataset.

#### Appendix J: Symbols

| Symbol | Name and description | Main units | Comments |
| --- | --- | --- | --- |
| $y$ | biomass yield: (rate of) biomass grown per (flux of) electron donor consumed | $\mathrm{Cmol\,mol^{-1}}$ | raw data available from references, defined according to SI B 2 |
| $\sigma$ | dissipation (per electron donor): free energy dissipated (entropy produced times temperature) per electron donor consumed | $\mathrm{kJ\,mol^{-1}}$ | computed according to Eq. (B22), (12) |
| $\bar{y}$ | mean yield within metabolic type | $\mathrm{Cmol\,mol^{-1}}$ | computed according to SI C |
| $\bar{\sigma}$ | mean dissipation within metabolic type | $\mathrm{kJ\,mol^{-1}}$ | computed according to SI C |
| $\delta y$ | yield deviation from $\bar{y}$ within metabolic type | $\mathrm{Cmol\,mol^{-1}}$ | computed according to SI C, SI E 2 |
| $\delta\sigma$ | dissipation deviation from $\bar{\sigma}$ within metabolic type | $\mathrm{kJ\,mol^{-1}}$ | computed according to SI C, SI E 2 |
| $\alpha$ | (dissipative) cost of growth: free energy dissipated (entropy produced times temperature) per biomass grown | $\mathrm{kJ\,Cmol^{-1}}$ | computed according to Eq. (6), SI C |
| $\beta$ | deviations coefficient within metabolic type | $\mathrm{kJ\,Cmol^{-1}}$ | obtained from linear regression according to Eq. (7), SI C |
| $\sigma_{\mathrm{sb/pd}}$ | free energy influx / outflux of substrates / products per electron donor consumed (with overbar is the mean within metabolic type) | $\mathrm{kJ\,mol^{-1}}$ | computed according to Eq. (B22) |
| $g/\rho_{\mathrm{C}}$ | free energy density of biomass $g$ per biomass carbon density $\rho_{\mathrm{C}}$ (with overbar is the mean within metabolic type) | $\mathrm{kJ\,Cmol^{-1}}$ | estimated from element composition according to SI H 2 |
| $r_{\mathrm{ed}}$ | "catabolic" force: thermodynamic force coupled to electron donor influx or yield (= 1) | $\mathrm{kJ\,mol^{-1}}$ | computed according to Eq. (E2), SI E 2 |
| $r_{\mathrm{b}}$ | "anabolic" force: thermodynamic force coupled to biomass growth rate or yield | $\mathrm{kJ\,Cmol^{-1}}$ | computed according to Eq. (E2), SI E 2 |
| $y_{i_{\mathrm{i}}}$ | independent yield: yield set as input data based on the original reference | $\mathrm{(C)mol\,mol^{-1}}$ | includes biomass, electron donor (= 1) and possibly others |
| $y_{i_{\mathrm{d}}}$ | dependent yield: yield determined by element conservation and independent yields | $\mathrm{mol\,mol^{-1}}$ | computed according to Eq. (E1),(4) |
| $\Delta$ | yield-force contribution to dissipation due to products/substrates independent yields | $\mathrm{kJ\,mol^{-1}}$ | = 0 when biomass and electron donor are the only independent variables; computed according to Eq. (E2),(12) |
| $\eta$ | (thermodynamic) efficiency: ratio of independent contributions to dissipation, free energy producing over free energy dissipating | | computed according to Eq. (E6) |

[16] Rob Phillips, Jane Kondev, Julie Theriot, and Hernan Garcia. *Physical biology of the cell*. Garland Science, 2012.

[17] Nick Lane and William Martin. The energetics of genome complexity. *Nature*, 467(7318):929–934, 2010.

[18] Michael Lynch and Georgi K Marinov. The bioenergetic costs of a gene. *Proceedings of the National Academy of Sciences*, 112(51):15690–15695, 2015.

[19] Gita Mahmoudabadi, Rob Phillips, Michael Lynch, and Ron Milo. Defining the energetic costs of cellular structures. *bioRxiv*, page 666040, 2019.

[20] Harold J Morowitz. Energy flow in biology; biological organization as a problem in thermal physics. 1968.

[21] TM McCollom and JP Amend. A thermodynamic assessment of energy requirements for biomass synthesis by chemolithoautotrophic micro-organisms in oxic and anoxic environments. *Geobiology*, 3(2):135–144, 2005.

[22] Jan P Amend, Douglas E LaRowe, Thomas M McCollom, and Everett L Shock. The energetics of organic synthesis inside and outside the cell. *Philosophical Transactions of the Royal Society B: Biological Sciences*, 368(1622):20120255, 2013.

[23] U Von Stockar and J-S Liu. Does microbial life always feed on negative entropy? thermodynamic analysis of microbial growth. *Biochimica et Biophysica Acta (BBA)-Bioenergetics*, 1412(3):191–211, 1999.

[24] Edwin H Battley. *Energetics of microbial growth*. Wiley, 1987.

[32] Edwin H Battley. Calculation of thermodynamic properties of protein in escherichia coli k-12 grown on succinic acid, energy

changes accompanying protein anabolism, and energetic role of atp in protein synthesis. *Biotechnology and bioengineering*, 40(2):280–288, 1992.

[33] Artur Wachtel, Riccardo Rao, and Massimiliano Esposito. Free-energy transduction in chemical reaction networks: From enzymes to metabolism. *The Journal of Chemical Physics*, 157(2), 2022.

[34] Sybren Ruurds De Groot and Peter Mazur. *Non-equilibrium thermodynamics*. Courier Corporation, 2013.

[62] Michiel Rutgers, Hanneke ML van der Gulden, and Karelvan Dam. Thermodynamic efficiency of bacterial growth calculated from growth yield of pseudomonas oxalaticus ox1 in the chemostat. *Biochimica et Biophysica Acta (BBA)-Bioenergetics*,

973(2):302–307, 1989.

[82] Philippe Duboc. Transient growth of saccharomyces cerevisiae. Technical report, EPFL, 1997.

[90] ANNE Belaich and JEAN-PIERRE Belaich. Microcalorimetric study of the anaerobic growth of escherichia coli: measure-

ments of the affinity of whole cells for various energy substrates. *Journal of bacteriology*, 125(1):19–24, 1976.
[91] Yasufumi Ishikawa, Yukio Nonoyama, and Makoto Shoda. Microcalorimetric study of aerobic growth of escherichia coli in batch culture. *Biotechnology and Bioengineering*, 23(12):2825–2836, 1981.
[92] Ian Marison and Urs Von Stockar. A calorimetric investigation of the aerobic cultivation of kluyveromyces fragilis on various substrates. *Enzyme and microbial technology*, 9(1):33–43, 1987.
[93] H Tamiya. Material and energy balances of biological synthesis. *Actualities Scientifiques et Industrielles*, 214, 1935.
[94] Syunzi Yamagata. *Über die elementare Zusammensetzung des Schimmelpilzkörpers*. 1934.
[95] Rita Whelton and Michael Doudoroff. Assimilation of glucose and related compounds by growing cultures of pseudomonas saccharophila. *Journal of Bacteriology*, 49(2):177–186, 1945.
[96] H Samejima and J Myers. On the heterotrophic growth of chlorella pyrenoidosa. *Microbiology*, 18(1):107–117, 1958.
[97] Sam R Hoover and Franklin E Allison. The growth metabolism of rhizobium, with evidence on the interrelations between respiration and synthesis. *Journal of Biological Chemistry*, 134(1):181–192, 1940.
[98] J-S Liu, IW Marison, and U Von Stockar. Microbial growth by a net heat up-take: a calorimetric and thermodynamic study on acetotrophic methanogenesis by methanosarcina barkeri. *Biotechnology and bioengineering*, 75(2):170–180, 2001.
[99] Rodrigo Patino, Marcel Janssen, and Urs von Stockar. A study of the growth for the microalga chlorella vulgaris by photo-bio-calorimetry and other on-line and off-line techniques. *Biotechnology and bioengineering*, 96(4):757–767, 2007.
[100] J Zhao, T Baba, H Mori, and K Shimizu. Global metabolic response of escherichia coli to gnd or zwf gene-knockout, based on 13 c-labeling experiments and the measurement of enzyme activities. *Applied microbiology and biotechnology*, 64: 91–98, 2004.
[101] Stanley E Mainzer and Walter P Hempfling. Effects of growth temperature on yield and maintenance during glucose-limited continuous culture of escherichia coli. *Journal of bacteriology*, 126(1):251–256, 1976.
[102] Cornelis Verduyn. Energetic aspects of metabolic fluxes in yeasts. 1992.
[103] Cornelis Verduyn, Erik Postma, W Alexander Scheffers, and Johannes P van Dijken. Physiology of saccharomyces cerevisiae in anaerobic glucose-limited chemostat culturesx. *Microbiology*, 136(3):395–403, 1990.
[104] Sébastien Bernacchi, Simon Rittmann, Arne H Seifert, Alexander Krajete, and Christoph Herwig. Experimental methods for screening parameters influencing the growth to product yield (y (x/ch4)) of a biological methane production (bmp) process performed with methanothermobacter marburgensis. *AIMS Bioengineering*, 1(2):72–87, 2014.
[105] Robert J Silbey, Robert A Alberty, George A Papadantonakis, and Moungi G Bawendi. *Physical chemistry*. John Wiley & Sons, 2022.
[106] Robert A Alberty. *Thermodynamics of biochemical reactions*. John Wiley & Sons, 2005.
[107] Moritz E Beber, Mattia G Gollub, Dana Mozaffari, Kevin M Shebek, Avi I Flamholz, Ron Milo, and Elad Noor. equilibrator 3.0: a database solution for thermodynamic constant estimation. *Nucleic acids research*, 50(D1):D603–D609, 2022.
[108] William R Cannon and Lionel M Raff. The formulation of chemical potentials and free energy changes in biochemical reactions. *Physical Chemistry Chemical Physics*, 23(27):14783–14795, 2021.
[109] Antonio Sabatini, Alberto Vacca, and Stefano Iotti. Balanced biochemical reactions: a new approach to unify chemical and biochemical thermodynamics. *Plos one*, 7(1):e29529, 2012.
[110] Elad Noor, Hulda S. Haraldsdóttir, Ron Milo, and Ronan M. T. Fleming. Consistent estimation of gibbs energy using component contributions. *PLOS Computational Biology*, 9(7):1–11, 07 2013. doi:10.1371/journal.pcbi.1003098.
[111] Edwin H. Battley and Handling Editor Daniel E. Dykhuizen. A theoretical study of the thermodynamics of microbial growth using saccharomyces cerevisiae and a different free energy equation. *The Quarterly Review of Biology*, 88(2):69–96, 2013. ISSN 00335770, 15397718.
[112] Robert W Sterner and James J Elser. Ecological stoichiometry. In *Ecological stoichiometry*. Princeton university press, 2017.
[113] T Gurakan, IW Marison, U Von Stockar, L Gustafsson, and E Gnaiger. Proposals for a standardized sample handling procedure for the determination of elemental composition and enthalpy of combustion of biological material. *Thermochimica acta*, 172:251–266, 1990.
[114] Marko Popovic. Thermodynamic properties of microorganisms: determination and analysis of enthalpy, entropy, and gibbs free energy of biomass, cells and colonies of 32 microorganism species. *Heliyon*, 5(6), 2019.
[115] Ron Grosz and Gregory Stephanopoulos. Statistical mechanical estimation of the free energy of formation of e. coli biomass for use with macroscopic bioreactor balances. *Biotechnology and bioengineering*, 25(9):2149–2163, 1983.
[116] Urs von Stockar, Lena Gustafsson, Christer Larsson, and Ian Marison. Thermodynamic considerations in constructing energy balances. *Biochimica et Biophysica Acta*, 1183:221–240, 1993.
[117] WM Thornton. Xv. the relation of oxygen to the heat of combustion of organic compounds. *The London, Edinburgh, and Dublin Philosophical Magazine and Journal of Science*, 33(194):196–203, 1917.
[118] Jean-Louis Cordier, Bertram M Butsch, Bernard Birou, and Uros von Stockar. The relationship between elemental composition and heat of combustion of microbial biomass. *Applied Microbiology and Biotechnology*, 25:305–312, 1987.
[119] Snehal A Patel and LE Erickson. Estimation of heats of combustion of biomass from elemental analysis using available electron concepts. *Biotechnology and Bioengineering*, 23(9):2051–2067, 1981.
[120] E Gnaiger and G Bitterlich. Proximate biochemical composition and caloric content calculated from elemental chn analysis: a stoichiometric concept. *Oecologia*, 62:289–298, 1984.
[121] Robert Hołyst and Andrzej Poniewierski. *Thermodynamics for chemists, physicists and engineers*, volume 344. Springer, 2012.

[122] Edwin H. Battley, Robert L. Putnam, and Juliana Boerio-Goates. Heat capacity measurements from 10 to 300 k and derived thermodynamic functions of lyophilized cells of saccharomyces cerevisiae including the absolute entropy and the entropy of formation at 298.15 k. *Thermochimica Acta*, 298(1):37–46, 1997. ISSN 0040-6031.

[123] Edwin H. Battley. An empirical method for estimating the entropy of formation and the absolute entropy of dried microbial biomass for use in studies on the thermodynamics of microbial growth. *Thermochimica Acta*, 326:7–15, 1999.

[124] Marko Popovic, Gavin B.G. Stenning, Axel Göttlein, and Mirjana Minceva. Elemental composition, heat capacity from 2 to 300 k and derived thermodynamic functions of 5 microorganism species. *Journal of Biotechnology*, 331:99–107, 2021. ISSN 0168-1656.

[125] Peter Linstrom and William Mallard. The nist chemistry webbook: A chemical data resource on the internet. (46), retrieved September 27, 2023.
